# Genome-wide association study provides new insights into the genetic architecture and pathogenesis of heart failure

**DOI:** 10.1101/682013

**Authors:** Sonia Shah, Albert Henry, Carolina Roselli, Honghuang Lin, Garðar Sveinbjörnsson, Ghazaleh Fatemifar, Åsa K. Hedman, Jemma B. Wilk, Michael P. Morley, Mark D. Chaffin, Anna Helgadottir, Niek Verweij, Abbas Dehghan, Peter Almgren, Charlotte Andersson, Krishna G. Aragam, Johan Ärnlöv, Joshua D. Backman, Mary L. Biggs, Heather L. Bloom, Jeffrey Brandimarto, Broad AF Investigators, Michael R. Brown, Leonard Buckbinder, David J. Carey, Regeneron Genetics Center, Daniel I. Chasman, Xing Chen, Xu Chen, Jonathan Chung, William Chutkow, James P. Cook, Graciela E. Delgado, Spiros Denaxas, Alexander S. Doney, Marcus Dörr, Samuel C. Dudley, Michael E. Dunn, EchoGen Consortium, Gunnar Engström, Tõnu Esko, Stephan B. Felix, Chris Finan, Ian Ford, Mohsen Ghanbari, Sahar Ghasemi, Vilmantas Giedraitis, Franco Giulianini, John S. Gottdiener, Stefan Gross, Daníel F. Guðbjartsson, Rebecca Gutmann, Christopher M. Haggerty, Pim van der Harst, Craig L. Hyde, Erik Ingelsson, J. Wouter Jukema, Maryam Kavousi, Kay-Tee Khaw, Marcus E. Kleber, Lars Køber, Andrea Koekemoer, Claudia Langenberg, Lars Lind, Cecilia M. Lindgren, Barry London, Luca A. Lotta, Ruth C. Lovering, Jian’an Luan, Patrik Magnusson, Anubha Mahajan, Kenneth B. Margulies, Winfried März, Olle Melander, Ify R. Mordi, Thomas Morgan, Andrew D. Morris, Andrew P. Morris, Alanna C. Morrison, Michael W. Nagle, Christopher P. Nelson, Alexander Niessner, Teemu Niiranen, Michelle L. O’Donoghue, Anjali T. Owens, Colin N. A. Palmer, Helen M. Parry, Markus Perola, Eliana Portilla-Fernandez, Bruce M. Psaty, Kenneth M. Rice, Paul M. Ridker, Simon P. R. Romaine, Jerome I. Rotter, Perttu Salo, Veikko Salomaa, Jessica van Setten, Alaa A. Shalaby, Diane T. Smelser, Nicholas L. Smith, Steen Stender, David J. Stott, Per Svensson, Mari-Liis Tammesoo, Kent D. Taylor, Maris Teder-Laving, Alexander Teumer, Guðmundur Thorgeirsson, Unnur Thorsteinsdottir, Christian Torp-Pedersen, Stella Trompet, Benoit Tyl, Andre G. Uitterlinden, Abirami Veluchamy, Uwe Völker, Adriaan A. Voors, Xiaosong Wang, Nicholas J. Wareham, Dawn Waterworth, Peter E. Weeke, Raul Weiss, Kerri L. Wiggins, Heming Xing, Laura M. Yerges-Armstrong, Bing Yu, Faiez Zannad, Jing Hua Zhao, Harry Hemingway, Nilesh J. Samani, John J.V. McMurray, Jian Yang, Peter M. Visscher, Christopher Newton-Cheh, Anders Malarstig, Hilma Holm, Steven A. Lubitz, Naveed Sattar, Michael V. Holmes, Thomas P. Cappola, Folkert Asselbergs, Aroon D. Hingorani, Karoline Kuchenbaecker, Patrick T. Ellinor, Chim C. Lang, Kari Stefansson, J. Gustav Smith, Ramachandran S. Vasan, Daniel I. Swerdlow, R. Thomas Lumbers

**Affiliations:** Institute for Molecular Bioscience, The University of Queensland, Brisbane, Queensland 4072, Australia; Institute of Cardiovascular Science, University College London, UK; Institute of Health Informatics, University College London, UK; Program in Medical and Population Genetics, The Broad Institute of MIT and Harvard, Cambridge, MA, USA; Department of Cardiology, University Medical Center Groningen, University of Groningen, Groningen, the Netherlands; Section of Computational Biomedicine, Department of Medicine, Boston University School of Medicine, Boston, Massachusetts, USA; National Heart, Lung, and Blood Institute’s and Boston University’s Framingham Heart Study, Framingham, Massachusetts, USA; deCODE genetics/Amgen Inc., Sturlugata 8, 101 Reykjavik, Iceland; Health Data Research UK London, University College London, UK; Cardiovascular Medicine unit, Department of Medicine Solna, Karolinska Institute, Stockholm, Sweden; Pfizer Worldwide Research & Development, 1 Portland St, Cambridge, MA, USA; Penn Cardiovascular Institute, Perelman School of Medicine, University of Pennsylvania, Philadelphia; Department of Epidemiology and Biostatistics, Imperial College London, St Mary’s Campus, London W2 1PG, UK; MRC-PHE Centre for Environment and Health, Department of Epidemiology and Biostatistics, Imperial College London, St Mary’s Campus, London W2 1PG, UK; Department of Clinical Sciences, Lund University, Malmö, Sweden; Department of Cardiology, Herlev Gentofte Hospital, Herlev Ringvej 57, 2650 Herlev, Denmark; Center for Genomic Medicine, Massachusetts General Hospital, Boston, MA, USA; Cardiovascular Research Center, Massachusetts General Hospital, Boston, MA, USA; Department of Neurobiology, Care Sciences and Society/Section of Family Medicine and Primary Care, Karolinska Institutet, Stockholm, Sweden; School of Health and Social Sciences, Dalarna University, Falun Sweden; Regeneron Genetics Center, 777 Old Saw Mill River Road, Tarrytown, NY 10591, USA; Regeneron Pharmaceuticals, Cardiovascular Research, 777 Old Saw Mill River Road, Tarrytown, NY 10591, USA; Department of Biostatistics, University of Washington, Seattle WA, USA; Department of Medicine, University of Washington, Seattle WA, USA; Division of Cardiology, Department of Medicine, Emory University Medical Center, Atlanta, GA, USA; Department of Epidemiology, Human Genetics, and Environmental Sciences, The University of Texas School of Public Health, Houston, Texas, USA; Department of Molecular and Functional Genomics, Geisinger; Danville, PA, USA; Division of Preventive Medicine, Brigham and Women’s Hospital, Boston, MA 02215; Harvard Medical School, Boston, MA 02115; Department of Medical Epidemiology and Biostatistics, Karolinska Institutet, Stockholm, Sweden; Novartis Institutes for Biomedical Research; Department of Biostatistics, University of Liverpool, Liverpool, UK; Vth Department of Medicine (Nephrology, Hypertensiology, Endocrinology, Diabetology, Rheumatology), Medical Faculty of Mannheim, University of Heidelberg, Germany; Division of Molecular & Clinical Medicine, University of Dundee, Ninewells Hospital and Medical School, Dundee DD1 9SY – Scotland, United Kingdom; Department of Internal Medicine B, University Medicine Greifswald, Greifswald, Germany; DZHK (German Center for Cardiovascular Research), partner site Greifswald, Greifswald, Germany; Cardiovascular Division, Department of Medicine, University of Minnesota, Minneapolis, MN, USA; Estonian Genome Center, Institute of Genomics, University of Tartu, Tartu, 51010, Estonia; Robertson Center for Biostatistics, University of Glasgow, United Kingdom; Department of Epidemiology, Erasmus University Medical Center, Rotterdam, the Netherlands; Institute for Community Medicine, University Medicine Greifswald, Greifswald, Germany; Department of Public Health and Caring Sciences, Geriatrics, Uppsala University, Uppsala 751 85, Sweden; Department of Medicine, Division of Cardiology, University of Maryland School of Medicine, Baltimore, MD; School of Engineering and Natural Sciences, University of Iceland, 101 Reykjavik, Iceland; Division of Cardiovascular Medicine, University of Iowa Carver College of Medicine, Iowa City, IA, USA; Department of Genetics, University Medical Center Groningen, University of Groningen, Groningen, The Netherlands; Durrer Center for Cardiogenetic Research, ICIN-Netherlands Heart Institute, Utrecht, The Netherlands; Department of Medicine, Division of Cardiovascular Medicine, Stanford University School of Medicine, Stanford, CA 94305; Stanford Cardiovascular Institute, Stanford University, Stanford, CA 94305; Department of Medical Sciences, Molecular Epidemiology and Science for Life Laboratory, Uppsala University, Uppsala, Sweden; Stanford Diabetes Research Center, Stanford University, Stanford, CA 94305; Department of Cardiology, Leiden University Medical Center, Leiden, the Netherlands; Einthoven Laboratory for Experimental Vascular Medicine, LUMC, Leiden, the Netherlands; Department of Public Health and Primary Care, University of Cambridge, Cambridge, CB2 0QQ, UK; Department of Cardiology, Copenhagen University Hospital Rigshospitalet, Copenhagen, Denmark; Department of Cardiovascular Sciences, University of Leicester and NIHR Leicester Biomedical Research Centre, Glenfield Hospital, Leicester; MRC Epidemiology Unit, Institute of Metabolic Science, University of Cambridge School of Clinical Medicine, Cambridge, CB2 0QQ, UK; Department of Medical Sciences, Uppsala University, Sweden; Big Data Institute at the Li Ka Shing Centre for Health Information and Discovery, University of Oxford, Oxford, UK; Wellcome Trust Centre for Human Genetics, University of Oxford, Oxford, UK; Division of Cardiovascular Medicine and Abboud Cardiovascular Research Center, University of Iowa, Iowa City, IA, USA; Synlab Academy, Synlab Holding Deutschland GmbH, Mannheim, Germany; Clinical Institute of Medical and Chemical Laboratory Diagnostics, Medical University of Graz, Graz, Austria; Department of Internal Medicine, Clinical Sciences, Lund University and Skåne University Hospital, Malmö, Sweden; Vanderbilt University School of Medicine; Usher Institute of Population Health Sciences and Informatics, University of Edinburgh, Edinburgh, United Kingdom; Department of Internal Medicine II, Division of Cardiology, Medical University of Vienna, Austria; National Institute for Health and Welfare, Helsinki, Finland; Department of Medicine, Turku University Hospital and University of Turku, Turku, Finland; TIMI Study Group, Cardiovascular Division, Brigham and Women’s Hospital, Boston, Massachusetts; Division of Vascular Medicine and Pharmacology, Department of Internal Medicine, Erasmus University Medical Center, Rotterdam, the Netherlands; Department of Medicine, Epidemiology, and Health Services, University of Washington, Seattle WA, USA; Kaiser Permanente Washington Health Research Institute, Kaiser Permanente Washington, Seattle WA, USA; The Institute for Translational Genomics and Population Sciences, Departments of Pediatrics and Medicine, Los Angeles Biomedical Research Institute at Harbor-UCLA Medical Center, Torrance, CA USA; Department of Cardiology, Division Heart and Lungs, University Medical Center Utrecht, University of Utrecht, Utrecht, The Netherlands; Division of Cardiology, Department of Medicine, University of Pittsburgh Medical Center and VA Pittsburgh HCS, Pittsburgh, PA, USA; Department of Epidemiology, University of Washington, Seattle WA, USA; Seattle Epidemiologic Research and Information Center, Department of Veterans Affairs Office of Research & Development, Seattle WA, USA; Department of Clinical Biochemistry, Copenhagen University Hospital, Herlev and Gentofte, Denmark; Institute of Cardiovascular and Medical Sciences, College of Medical, Veterinary and Life Sciences, University of Glasgow, United Kingdom; Department of Clinical Science and Education, Södersjukhuset, Karolinska Institutet, Stockholm, Sweden; Department of Cardiology, Södersjukhuset, Stockholm, Sweden; Institute for Translational Genomics and Population Sciences, LABiomed and Departments of Pediatrics at Harbor-UCLA Medical Center, Torrance, Calif., USA 90502; Division of Cardiology, Department of Internal Medicine, Landspitali, National University Hospital of Iceland, Hringbraut, 101 Reykjavik, Iceland; Faculty of Medicine, Department of Medicine, University of Iceland, Saemundargata 2, 101 Reykjavik, Iceland; Department of Epidemiology and Biostatistics, Aalborg University Hospital, Aalborg, Denmark; Department of Health, Science and Technology, Aalborg University Hospital, Denmark; Departments of Cardiology, Aalborg University Hospital, Denmark; Section of Gerontology and Geriatrics, Department of Internal Medicine, Leiden University Medical Center, Leiden, the Netherlands; Translational and Clinical Research, Servier Cardiovascular Center for Therapeutic Innovation, 50 rue Carnot 92284 Suresnes, France; Department of Internal Medicine, Erasmus MC, University Medical Center Rotterdam, Rotterdam, the Netherlands; Interfaculty Institute for Genetics and Functional Genomics, University Medicine Greifswald, Greifswald, Germany; Human Genetics, GlaxoSmithKline, Collegeville, PA; Division of Cardiovascular Medicine, Department of Internal Medicine, The Ohio State University Medical Center, Columbus, OH, USA; Université de Lorraine, CHU de Nancy, Inserm and INI-CRCT (F-CRIN), Institut Lorrain du Coeur et des Vaisseaux, 54500 Vandoeuvre Lès Nancy, France; The National Institute for Health Research University College London Hospitals Biomedical Research Centre, University College London; BHF Cardiovascular Research Centre, University of Glasgow, Scotland, United Kingdom; Queensland Brain Institute, The University of Queensland, Brisbane, Queensland 4072, Australia; Center for Human Genetic Research, Massachusetts General Hospital, Boston, MA, USA; Cardiac Arrhythmia Service and Cardiovascular Research Center, Massachusetts General Hospital, Boston, MA, USA; Medical Research Council Population Health Research Unit at the University of Oxford, Oxford, UK; Clinical Trial Service Unit and Epidemiological Studies Unit, Nuffield Department of Population Health, Big Data Institute, University of Oxford, Oxford, UK; National Institute for Health Research Oxford Biomedical Research Centre, Oxford University Hospital, Oxford, UK; Division of Psychiatry, University College of London, London W1T 7NF, UK; UCL Genetics Institute, University College London, London WC1E 6BT, UK; Department of Cardiology, Clinical Sciences, Lund University and Skåne University Hospital, Lund, Sweden; Wallenberg Center for Molecular Medicine and Lund University Diabetes Center, Lund University, Lund, Sweden; Sections of Cardiology, Preventive Medicine and Epidemiology, Department of Medicine, Boston University Schools of Medicine and Public Health, Boston, Massachusetts, USA; BenevolentAI, London, UK; Bart’s Heart Centre, St. Bartholomew’s Hospital, London, UK

## Abstract

Heart failure (HF) is a leading cause of morbidity and mortality worldwide^1^. A small proportion of HF cases are attributable to monogenic cardiomyopathies and existing genome-wide association studies (GWAS) have yielded only limited insights, leaving the observed heritability of HF largely unexplained^2–4^. We report the largest GWAS meta-analysis of HF to-date, comprising 47,309 cases and 930,014 controls. We identify 12 independent variant associations with HF at 11 genomic loci, all of which demonstrate one or more associations with coronary artery disease (CAD), atrial fibrillation, or reduced left ventricular function suggesting shared genetic aetiology. Expression quantitative trait analysis of non-CAD-associated loci implicate genes involved in cardiac development (*MYOZ1, SYNPO2L*), protein homeostasis (*BAG3*), and cellular senescence (*CDKN1A*). Using Mendelian randomisation analysis we provide new evidence supporting previously equivocal causal roles for several HF risk factors identified in observational studies, and demonstrate CAD-independent effects for atrial fibrillation, body mass index, hypertension and triglycerides. These findings extend our knowledge of the genes and pathways underlying HF and may inform the development of new therapeutic approaches.

## Introduction

Heart failure (HF) affects more than 30 million individuals worldwide and its prevalence is rising^1^. HF-associated morbidity and mortality remain high despite therapeutic advances, with five-year survival averaging ~50%^5^. HF is a clinical syndrome defined by fluid congestion and exercise intolerance due to cardiac dysfunction^6^. HF results typically from myocardial disease with impairment of left ventricular (LV) function manifesting with either reduced or preserved ejection fraction. Several cardiovascular and systemic disorders are implicated as aetiological factors, most notably coronary artery disease (CAD), obesity and hypertension; multiple risk factors frequently cooccur and the contribution to aetiology has been challenging based on observational data alone^7,8^. Monogenic hypertrophic and dilated cardiomyopathy (DCM) syndromes are known causes of HF, although they account for a small proportion of disease burden^9^. HF is a complex disorder with an estimated heritability of ~26%^2^. Previous modest-sized genome-wide association studies (GWAS) of HF reported two loci, while studies of DCM have identified a few replicated loci^10–14^. We hypothesised that a GWAS of HF with greater power would provide an opportunity for: (i) discovery of genetic variants modifying disease susceptibility in a range of comorbid contexts, both through subtype-specific and shared pathophysiological mechanisms, such as fluid congestion; and (ii) provide insights into aetiology by estimating the unconfounded causal contribution of observationally associated risk factors by Mendelian randomisation analysis^15^.

## Results

We conducted a large-scale GWAS comprising 47,309 cases and 930,014 controls of European ancestry across 26 studies from the The Heart Failure Molecular Epidemiology for Therapeutic Targets (HERMES) Consortium. Genotype data were imputed to either the 1000 Genomes Project (60%), Haplotype Reference Consortium (35%) or study-specific reference panels (5%). We performed a fixed-effect inverse variance-weighted meta-analysis relating 8,281,262 common and low-frequency variants (minor allele frequency > 1%) to HF risk (**Figure 1**). We identified 12 independent genetic variants, at 11 loci associated with HF at genome-wide significance (*P* < 5 × 10^−8^), including 10 loci not previously reported for heart failure (**Figure 2, Table 1**). The quantile-quantile, regional association plots and study-specific effects for each independent variant are shown in **Supplementary Figures 1-3**. We replicated two previously-reported associations for HF and three of four loci for DCM (Bonferroni-corrected *P* < 0.05) (**Supplementary Table 1**). Using linkage disequilibrium score regression (LDSC)^16^, we estimated the heritability of HF in UK Biobank (*h^2^_g_*) on the liability scale, as 0.088 (standard error (SE) = 0.013), based on an estimated disease prevalence of 2.5%^17^.

**Figure 1.**
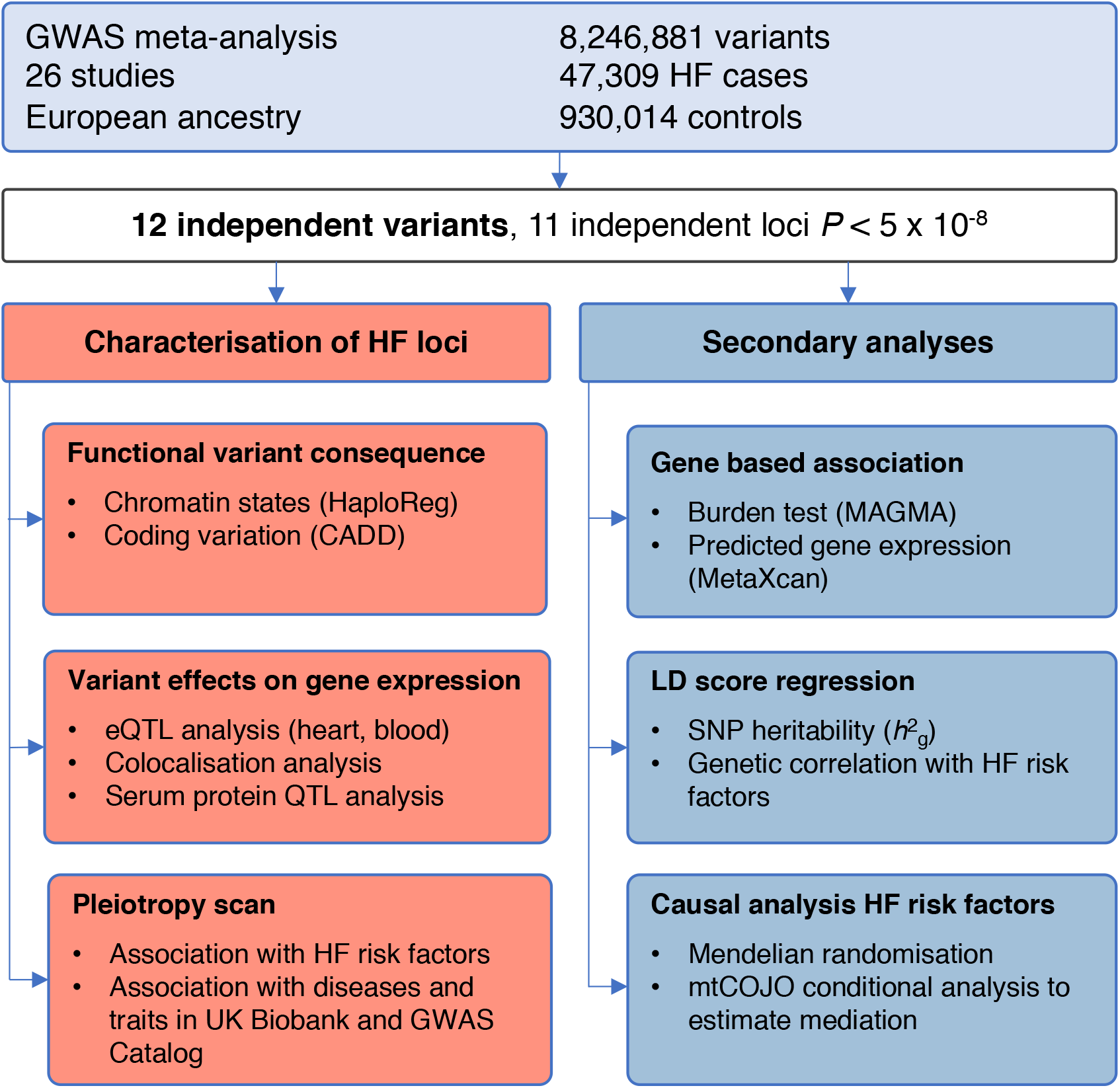
Study design and analysis workflow. Overview of study design to identify and characterise heart failure associated risk loci and for secondary cross-trait genome-wide analyses. Abbreviations: GWAS, genome-wide association study; QTL, quantitative trait locus; MAGMA, Multi-marker Analysis of GenoMic Annotation; SNP, single nucleotide polymorphism; mtCOJO, multi-trait-based conditional and joint analysis.

**Figure 2.**
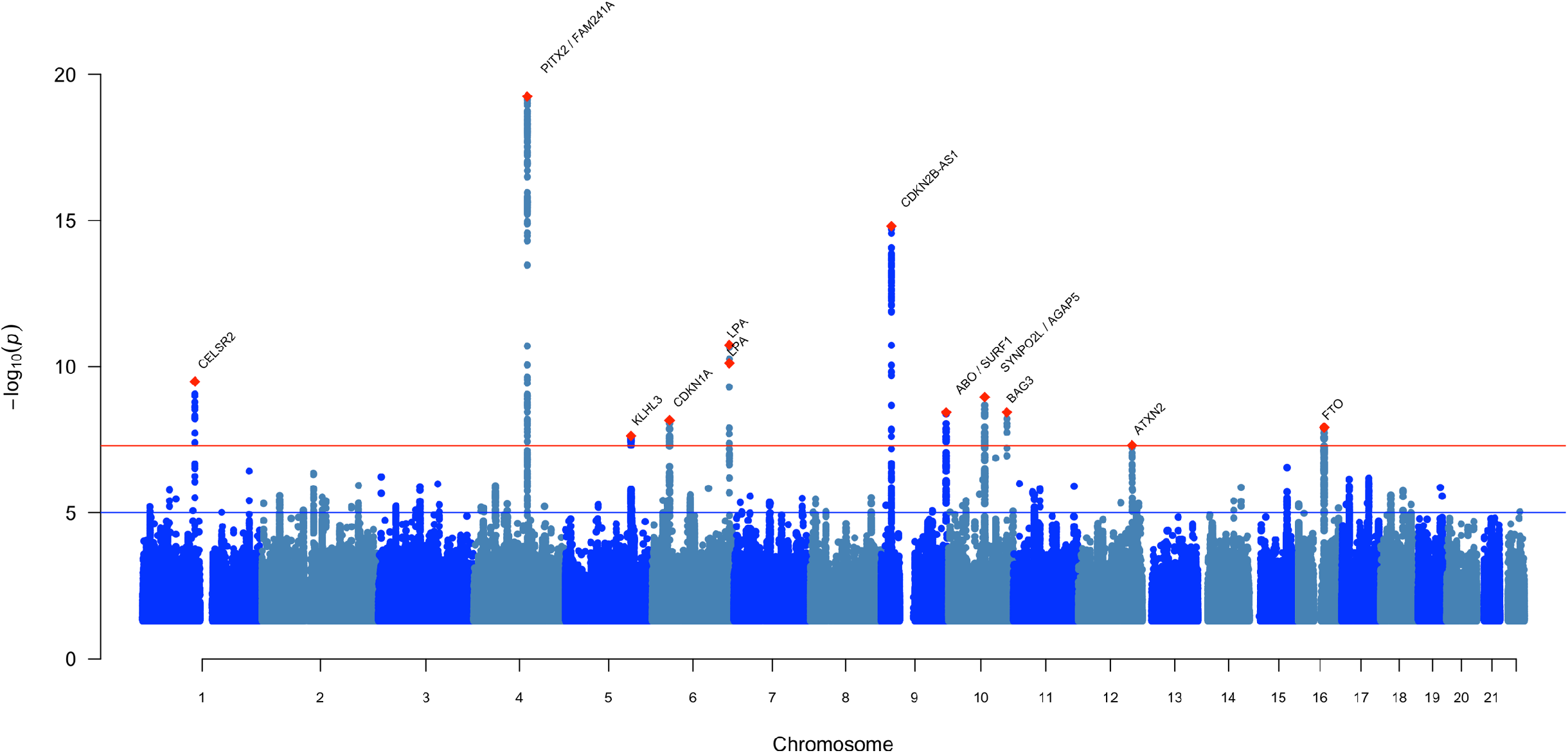
Manhattan plot of genome-wide heart failure associations. The x-axis represents the genome in physical order; the y-axis shows −log_10_ *P* values for individual variant association with heart failure risk from the meta-analysis (n = 977,323). Suggestive associations at a significance level of *P* < 1 × 10^−5^ are indicated by the blue line, while genome-wide significance at *P* < 5 × 10^−8^ is indicated by the red line. Meta-analysis was performed using a fixed-effect inverse-variance weighted model. Independent genome-wide significant variants are annotated with the nearest gene(s).

**Table 1.** Variants associated with heart failure at genome-wide significance

| rsID | Chr | Position (hg19) | Nearest gene(s)<sup>a</sup> | Function | Risk/ ref allele | RAF (%) | OR (95% CI) | P-value | I<sup>2</sup><sub>HET</sub> | P<sub>HET</sub> |
| --- | --- | --- | --- | --- | --- | --- | --- | --- | --- | --- |
| rs660240 | 1 | 109817838 | CELSR2 | UTR3 | C / T | 0.79 | 1.06 (1.04-1.08) | 3.25E-10 | 0 | 0.513 |
| rs17042102 | 4 | 111668626 | PITX2, FAM241A | intergenic | A / G | 0.12 | 1.12 (1.09-1.14) | 5.71E-20 | 43.1 | 0.008 |
| rs11745324 | 5 | 137012171 | KLHL3 | intronic | G / A | 0.77 | 1.05 (1.03-1.07) | 2.35E-08 | 5.7 | 0.381 |
| rs4135240 | 6 | 36647680 | CDKN1A | intronic | T / C | 0.66 | 1.05 (1.03-1.07) | 6.84E-09 | 43.8 | 0.009 |
| rs55730499 | 6 | 161005610 | LPA | intronic | T / C | 0.07 | 1.11 (1.08-1.14) | 1.83E-11 | 21.1 | 0.164 |
| rs140570886 | 6 | 161013013 | LPA | intronic | C / T | 0.02 | 1.24 (1.16-1.3) | 7.69E-11 | 24.8 | 0.133 |
| rs1556516 | 9 | 22100176 | CDKN2B-AS1 | ncRNA_intronic | C / G | 0.48 | 1.06 (1.05-1.08) | 1.57E-15 | 12.8 | 0.269 |
| rs600038 | 9 | 136151806 | ABO, SURF1 | intergenic | C / T | 0.21 | 1.06 (1.04-1.08) | 3.68E-09 | 0 | 0.729 |
| rs4746140 | 10 | 75417249 | SYNPO2L, AGAP5 | intergenic | G / C | 0.85 | 1.07 (1.05-1.09) | 1.10E-09 | 9.7 | 0.319 |
| rs17617337 | 10 | 121426884 | BAG3 | intronic | C / T | 0.78 | 1.06 (1.04-1.08) | 3.65E-09 | 55 | 0.000 |
| rs4766578 | 12 | 111904371 | ATXN2 | intronic | T / A | 0.47 | 1.04 (1.03-1.06) | 4.90E-08 | 10.6 | 0.308 |
| rs56094641 | 16 | 53806453 | FTO | intronic | G / A | 0.42 | 1.05 (1.03-1.06) | 1.21E-08 | 17.4 | 0.215 |

The table shows the 12 independent variants associated with HF at the genome-wide significance level (association $P$ -value $< 5 \times 10^{-8}$ ) in the meta-analysis of 29 studies. Meta-analyses were carried out using an inverse variance weighted fixed-effect approach. The $I^2_{\text{HET}}$ describes the percentage of variation across the 29 studies that is due to heterogeneity. $P_{\text{HET}}$ was derived from a Cochran's Q-test (two-sided) for heterogeneity. <sup>a</sup>Nearest gene with a functional protein or RNA (e.g. anti-sense RNA) product that either overlaps with the sentinel variant, or for intergenic variants, the nearest genes up- and downstream, respectively (separated by comma). Abbreviations: Chr, chromosome; ref, reference; RAF, risk allele frequency; OR, odds ratio; CI, confidence intervals; HET, heterogeneity; $I^2$ , I-squared.

Next, we investigated associations between the identified loci and other traits that may provide insights into aetiology. First, we queried the NHGRI-EBI GWAS Catalog^18^ and a large database of genetic associations in UK Biobank (http://www.nealelab.is/uk-biobank), and identified several biomarker and disease associations at each locus (**Supplementary Tables 2-3**). Second, we tested for associations of identified loci with ten known HF risk factors, including cardiac structure and function measures, using GWAS summary data (**Supplementary Table 4**)^19–26^. Six sentinel variants were also associated with CAD, including established loci such as *CDKN2B-AS1* (9p21) and *LPA*^21^. Four variants were associated with atrial fibrillation (AF), a common antecedent and sequela of HF^27^. To estimate whether the HF risk effects were mediated wholly or in part by risk factors upstream of HF (e.g. CAD), we conditioned HF GWAS summary statistics on nine HF risk factors using mtCOJO^28^ (**Supplementary Table 5**). Conditioning on AF attenuated the HF risk effect by > 50% for the *PITX2 / FAM241A* locus but not other AF-associated loci (*KLHL3, SYNPOL2 / AGAP5*), conditioning on CAD fully attenuated effects for two of the six CAD loci (*LPA, CDKN2B-AS1*) and conditioning on BMI ablated the effect of the *FTO* locus (**Supplementary Figure 4 and Supplementary Table 5**). Next we performed hierarchical agglomerative clustering of loci based on cross-trait associations to identify groups related to HF subtypes (**Figure 3**). Among HF loci not associated with CAD, a group of four clustered together, of which two (*KLHL3* and *SYNPO2L / AGAP5*) were associated with AF and two (*BAG3* and *CDKN1A*) with reduced LV systolic function (fractional shortening) (Bonferroni-corrected *P* < 0.05); we highlight the results for these loci in our reporting of subsequent analyses to identify candidate genes. Notably, genetic associations with DCM at the *BAG3* locus have been reported previously^13,14^.

**Figure 3.**
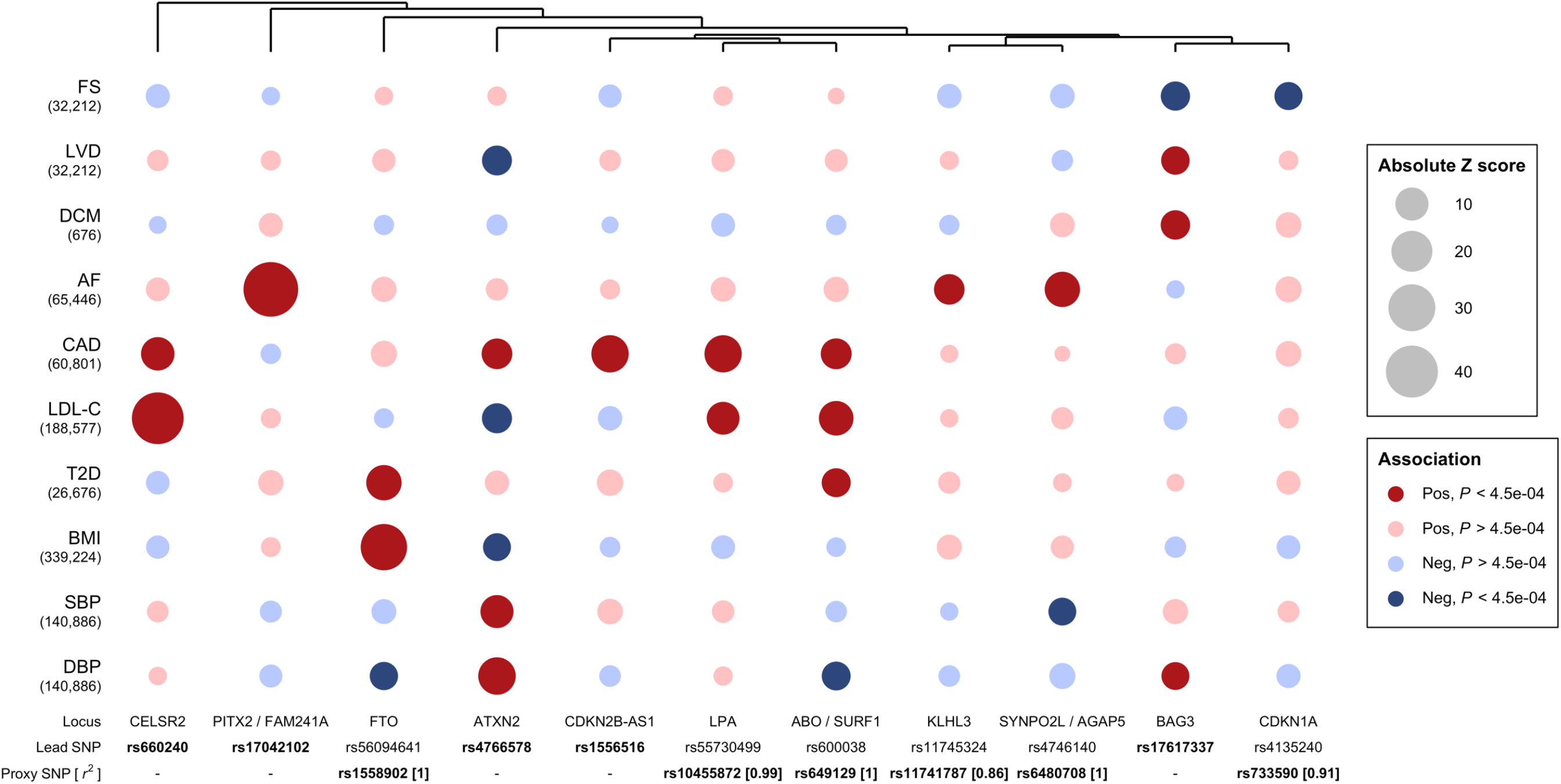
Associations of HF risk variants with traits relating to disease subtypes and risk factors. This bubble plot shows associations between the identified HF loci and risk factors and quantitative imaging traits, using summary estimates from UK Biobank (DCM, dilated cardiomyopathy) and published GWAS summary statistics. Number in bracket represents sample size (for quantitative traits) or number of cases (for binary traits) used to derive the GWAS summary statistics. The size of the bubble represents the absolute Z score for each trait, with the direction oriented towards the HF risk allele. Red / blue indicates a positive / negative cross-trait association (i.e. increase / decrease in disease risk or increase / decrease in continuous trait). We accounted for family-wise error rate at 0.05 by Bonferroni correction for the 10 traits tested per HF locus (*P* < 4.5e^−4^); traits meeting this threshold of significance for association are indicated by dark colour shading. Agglomerative hierarchical clustering of variants was performed using the complete linkage method, based on Euclidian distance. Where a sentinel variant was not available for all traits, a common proxy was selected (bold text). For the *LPA* locus, associations for the more common of the two variants at this locus are shown. Bold text represents variants whose estimates are plotted, upon which we performed hierarchical agglomerative clustering using the complete linkage method based on Euclidian distance. Abbreviations: FS, fractional shortening; LVD, left ventricular dimension; DCM, dilated cardiomyopathy; AF, atrial fibrillation; CAD, coronary artery disease; LDL-C, low density lipoprotein cholesterol; T2D, type 2 diabetes; BMI, body mass index; SBP, systolic blood pressure; DBP, diastolic blood pressure.

We performed gene-based association analyses using MAGMA^29^ to identify tissues and aetiological pathways relevant to HF. Thirteen genes were associated with HF at genome-wide significance, of which four were located within 1Mb of a sentinel HF variant and expressed in heart tissue (**Supplementary Table 6**). Tissue specificity analysis across 53 tissue types from the Genotype Tissue Expression (GTEx) project identified the atrial appendage as the highest ranked tissue for gene expression enrichment excluding reproductive organs (**Supplementary Figure 5**). We sought to map candidate genes to the HF loci by assessing the functional consequences of sentinel variants (or their proxies) on gene expression, and protein structure/abundance using quantitative trait locus (QTL) analyses. Given enrichment for gene expression of HF-associated genes in the atrial appendage, we prioritised heart tissues for mapping studies, including QTL analysis.

Since the identified HF variants were located in non-coding regions, we investigated if sentinel variants were in linkage disequilibrium (LD, *r*^2^ > 0.8) with non-synonymous variants with predicted deleterious effects. We identified a missense variant in *BAG3* (rs2234962; *r*^2^ = 0.99 with sentinel variant rs17617337) associated previously with DCM and progression to HF, and three missense variants in *SYNPO2L* (rs34163229, rs3812629, rs60632610; all *r*^2^ > 0.9 with sentinel variant rs4746140)^13,14,30^. All 4 missense variants had Combined Annotation Dependent Depletion scores >20 suggesting deleterious effects (**Supplementary Table 7**).

Given the enrichment of genes expressed in heart tissue from the MAGMA analysis, we examined the effects of HF variants on *cis* gene expression in LV, left atrium and right atrium auricular region tissues, using data from the Myocardial Applied Genomics Network (MAGNet) and Genotype-Tissue Expression (GTEx) project. Three of 12 variants were significantly associated with the expression of one or more genes located in *cis* in at least one heart tissue (Bonferroni-corrected *P* < 0.05) (**Supplementary Table 8**). Given the modest sample sizes of myocardial eQTL studies, we also investigated a large whole blood eQTL dataset (eQTLGen consortium, n = 31,684) and found associations with *cis* gene expression (*P* < 5×10^−8^) for 8 of 12 sentinel variants (**Supplementary Table 9**)^31^. For most HF variants, heart eQTL associations were consistent with those for blood traits; however, for intronic HF sentinel variants in *BAG3*, *CDKN1A* and *KLHL3* we detected expression of the corresponding gene transcripts in blood only.

Next, to prioritise among candidate genes identified through eQTL associations, we estimated the posterior probability for a common causal variant underlying associations with gene expression and HF at each locus, by conducting pairwise Bayesian colocalization analysis ^32^. We found evidence for colocalization (posterior probability > 0.7) for *MYOZ1* and *SYNPO2L* in heart, *PSRC1* and *ABO* in heart and blood; and *CDKN1A* in blood (**Supplementary Tables 8 and 9**). *PSRC1* and *MYOZ1* were also implicated in a transcriptome-wide association analysis performed using predicted gene expression based on GTEx human atrial and ventricular expression reference data (**Supplementary Table 10**). Using serum protein QTL data from the INTERVAL study (N = 3,301), we also identified significant concordant *cis* associations for *BAG3* and *ABO* (**Supplementary Table 11**)^33^.

The evidence linking candidate genes with HF risk loci is summarised in **Supplementary Table 12**, and candidate genes are described in the **Supplementary Note**. At HF risk loci associated with reduced systolic function of AF, but not with CAD, the annotated functions of candidate genes related to myocardial disease processes, and traits that may influence clinical expressivity, such as renal sodium handling. For example, the sentinel variant at the *SYNPO2L / AGAP5* locus was associated with expression of *MYOZ1* and *SYNPO2L*, encoding two α-actinin binding Z-disk cardiac proteins. *MYOZ1* is a negative regulator of calcineurin signalling, a pathway linked to pathological hypertrophy^34,35^ and *SYNPO2L* is implicated in cardiac development and sarcomere maintenance^36^.

The HF sentinel variant at the *BAG3* locus was in high LD with a non-synonymous variant associated previously with DCM,^14^ and was associated with decreased *cis* gene expression in blood. *BAG3* encodes a Z-disc-associated protein that mediates selective macroautophagy and promotes cell survival through interaction with apoptosis regulator *BCL2*^37^. *CDKN1A* encodes p21, a potent cell cycle inhibitor that mediates post-natal cardiomyocyte cell-cycle arrest^38^ and is implicated in LMNA-mediated cellular stress responses^39^. *KLHL3* is a negative regulator of the thiazide-sensitive Na^+^Cl^−^ cotransporter (*SLC12A3*) in the distal nephron; loss of function variants cause familial hyperkalaemic hypertension (FHHt) by increasing constitutive sodium and chloride resorption^40^. The sentinel variant at this locus was associated with decreased gene expression and could predispose to sodium and fluid retention. Notably, thiazide diuretics inhibit *SLC12A3* to restore sodium and potassium homeostasis in FHHt and are effective treatments for preventing hypertensive HF ^41^.

Although many risk factors are associated with HF, only myocardial infarction and hypertension have an established causal role in randomised controlled trials (RCTs)^42^. Important questions remain about causality for other risk factors. For instance, type 2 diabetes (T2D) is a risk factor for HF, yet it is unclear if the association is mediated via CAD risk or by direct myocardial effects, which may have important preventative implications^43^. Accordingly, we investigated potential causal roles for modifiable HF risk factors using GWAS summary data. First, we estimated the genetic correlation (r_*g*_) between HF and 11 related traits using bivariate LDSC. For eight of the eleven traits tested, we found evidence of shared additive genetic effects with estimates of r_*g*_ ranging from −0.25 to 0.67 (**Supplementary Table 13).** The estimated CAD-HF r_*g*_ was 0.67, suggesting 45% (*r_g_*^2^) of variation in genetic risk of HF is accounted for by common genetic variation shared with CAD, and that the remaining genetic variation is independent of CAD.

Next, we estimated the causal effects of the eleven HF risk factors using Generalised Summary-data-based Mendelian Randomisation, which accounts for pleiotropy by excluding heterogenous variants based on the HEIDI test (**Online Methods, Supplementary Figure 6, Supplementary Table 14**). Consistent with evidence from RCTs, we found evidence for causal effects of higher diastolic blood pressure (OR = 1.30 per 10 mmHg, *P* = 9.13 × 10^−21^) and systolic blood pressure (OR = 1.18 per 10 mmHg, *P* = 4.8 × 10^−23^), and higher risk of CAD (OR = 1.36, *P* = 1.67 × 10^−70^) on HF. We found a standard deviation increment of body mass index (BMI) (equivalent to 4.4 kg/m^2^ [men] – 5.4 kg/m^2^ [women]^44^) accounted for a 74% higher HF risk (*P* = 2.67 × 10^−50^), consistent with previous reports ^45,46^. We identified evidence supporting causal effects of genetic liability to AF (OR of HF per 1 log odds higher AF = 1.19, *P* = 1.40 × 10^−75^) and T2D (OR of HF per 1 log odds higher T2D = 1.05, *P* = 6.35 × 10^−05^) and risk of HF. We did not find supportive evidence for a causal role for higher heart rate or lower glomerular filtration rate despite reported observational associations^47,48^.

To investigate whether risk factor effects on HF were mediated by CAD and AF, we performed analyses conditioning for CAD and AF using mtCOJO. We observed attenuation of the effect of T2D after conditioning for CAD (OR = 1.02, *P* = 0.19), suggesting at least partial mediation by CAD risk rather than through direct myocardial effects of hyperglycaemia. Similarly, the effects of low density lipoprotein cholesterol (LDL-C) were fully explained by effects of CAD on HF risk (OR = 1.00, *P* = 0.80). Conversely, the effects of blood pressure, BMI, and triglycerides were only partially attenuated, suggesting causal mechanisms independent of those associated with AF and CAD (**Figure 4**, **Supplementary Table 14)**.

**Figure 4.**
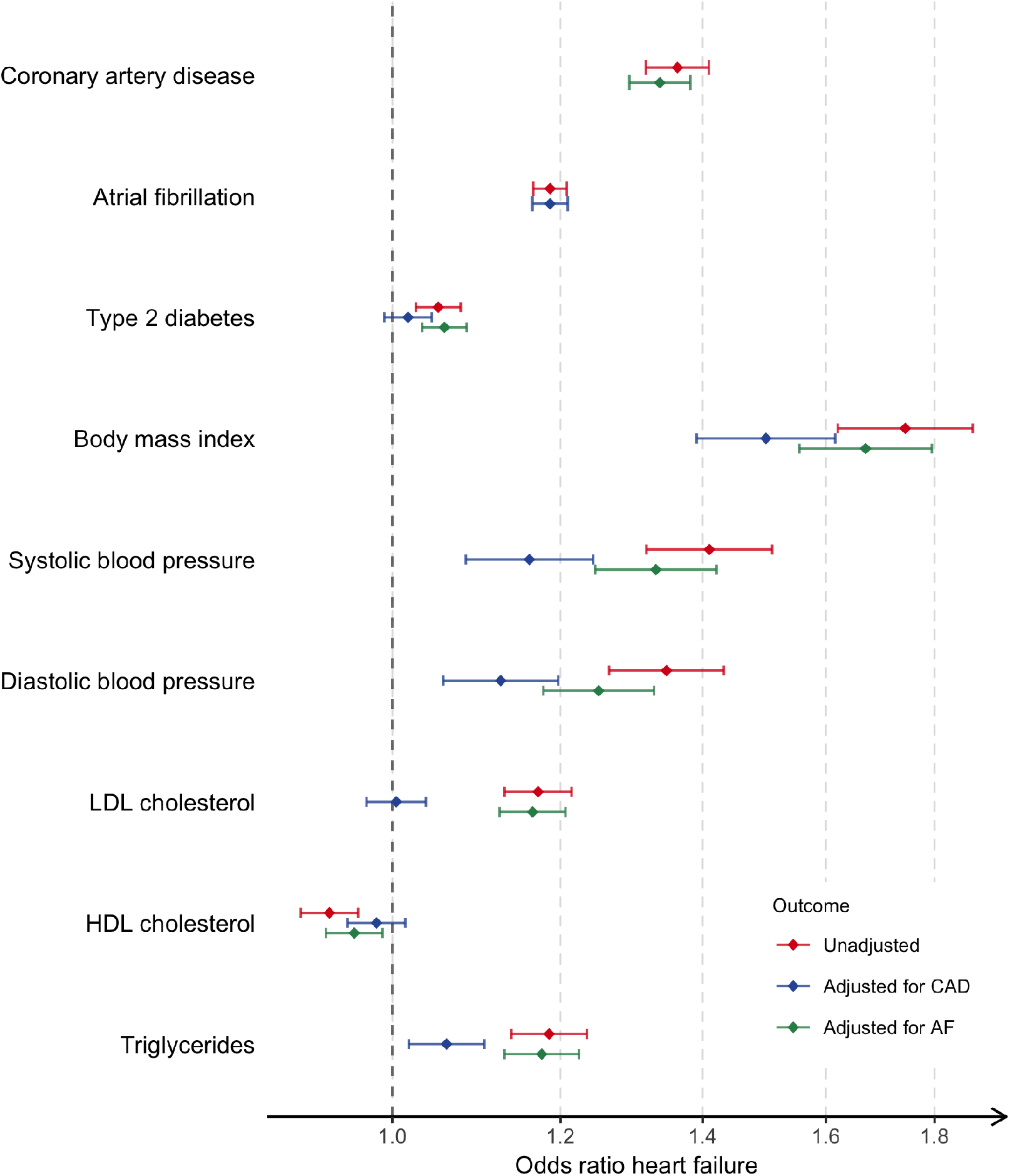
Conditional Mendelian randomisation analyses of HF risk factors. Forest plot of HF risk factors with significant causal effect HF risk estimated using Mendelian randomisation, implemented with GSMR. Diamonds represent the odds ratio and the error bars indicate the 95% confidence interval. The unadjusted estimates represent the risk of HF as estimated from the HF GWAS data, while the adjusted estimates represent risk of HF conditioned using GWAS summary statistics for atrial fibrillation (Adjusted for AF) or coronary artery disease (Adjusted for CAD) estimated using the mtCOJO method. For binary traits (coronary artery disease, atrial fibrillation, type 2 diabetes), the MR estimates represent average causal effect per natural-log odds increase in the trait risk. For continuous traits, the MR estimates represent average causal effect per standard deviation increase in the reported unit of the trait. Abbreviations: LDL, low density lipoprotein; HDL, high density lipoprotein; CAD, coronary artery disease; AF, atrial fibrillation.

We identify 12 independent variant associations for HF risk at 11 genomic loci by leveraging genome-wide data on 47,309 cases and 930,014 controls, including 10 loci not previously associated with HF. The identified loci were associated with modifiable risk factors and traits related to LV structure and function. Conditioning for CAD, AF and blood pressure traits demonstrated that the effects of some loci (e.g. CDKN2B-AS1 or 9p21) were mediated wholly via risk factor trait associations (e.g. CAD); however, for 8 of 12 variants the attenuation of effects was <50% suggesting alternative mechanisms may be important. Those loci associated with reduced LV systolic function or AF mapped to candidate genes implicated in processes of cardiac development, protein homeostasis, and cellular senescence. We use genetic causal inference and conditional analysis to explore the syndromic heterogeneity and causal biology of HF and provide new insights into aetiology. Mendelian randomisation analysis confirms previously reported casual effects for BMI and provides new evidence supporting the causal role of several observationally-linked risk factors including AF, elevated blood pressure (DBP and SBP), LDL-C, CAD, triglycerides, and T2D. Using conditional analysis we demonstrate CAD-independent effects for AF, BMI, blood pressure, and estimate that the effects of T2D are mostly mediated by an increased risk of CAD.

The heterogeneity of aetiology and clinical manifestation of heart failure are likely to have reduced statistical power. We identify a modest number of genetic associations for HF compared to other cardiovascular disease GWAS of comparable sample size, such as for AF, suggesting that an important component of HF heritability may be more attributable to specific disease subtypes than components of a final common pathway^49^. Subsequent studies will explore emerging opportunities to define HF subtypes and longitudinal phenotypes in large biobanks and patient registries at scale using standardised definitions based on diagnostic codes, imaging and electronic health records. We speculate that future analysis of HF subtypes may yield additional insights into the genetic architecture of HF to inform new approaches to prevention and treatment.

## Acknowledgements

A full list of acknowledgements is given in the Supplementary Note.

## Online Methods

### Samples

Participants of European ancestry from 26 cohorts (with a total of 29 distinct datasets) with either a case-control or population-based study design were included in the meta-analysis, as part of the Heart Failure Molecular Epidemiology for Therapeutic Targets (HERMES) Consortium. Cases included participants with a clinical diagnosis of heart failure of any aetiology with no inclusion criteria based on left ventricular ejection fraction; controls were participants without HF. Definitions used to adjudicate heart failure status within each study are detailed in the **Supplementary Table 15** and baseline characteristics for each study are provided in **Supplementary Table 16**. We meta-analysed data from a total of 47,309 cases and 930,014 controls. All included studies were ethically approved by local institutional review boards and all participants provided written informed consent.

### Genotyping and imputation

All studies used high-density genotyping arrays and performed genotype calling and pre-imputation quality control (QC) as reported in **Supplementary Table 17**. Studies performed imputation using one or more of the following reference panels: 1000 Genomes (Phase 1 or Phase 3) ^1^, Hapmap 2 NCBI build 36^2^, Haplotype Reference Consortium (HRC)^3^, the Estonian Whole Genome Sequence reference^4^, or a reference sample based on 15,220 whole genome sequences of Icelandic individuals. The following software tools were used by studies for phasing: Eagle^5^, MaCH^6^, SHAPEIT^7^; and imputation: mimimac2^8^, IMPUTE2^9^. For imputation to the HRC reference panel, the Sanger Imputation Server (https://www.sanger.ac.uk/science/tools/sanger-imputation-service) was used. The deCODE study was imputed using study specific procedures^10^. Methods for phasing, imputation and postimputation QC for each study are detailed in **Supplementary Table 17**.

### Study-level genome wide association analysis

Genome-wide association (GWA) analysis for each study was performed locally according to a common analysis plan, and summary level estimates were provided for meta-analysis. Autosomal single nucleotide polymorphisms were tested for association with HF using logistic regression, assuming additive genetic effects. For the Cardiovascular Health Study, HF association estimates were generated by analysis of incident cases using a Cox proportional hazards model. All studies included age and sex (except for single sex studies) as covariates in the regression models. Principal components (PCs) were included as covariates for individual studies as appropriate. The following tools were used for study-level GWA analysis: ProbABEL^11^, mach2dat (http://www.unc.edu/~yunmli/software.html), QuickTest^12^, PLINK2^13^, SNPTEST^14^, or R^15^ as detailed in **Supplementary Table 17**.

### Quality control on study summary-level data

Quality control of summary-level results for each study was performed according to the protocol described in Winkler et al.^16^. In brief, we used the EasyQC tool to harmonise variant IDs and alleles across studies and to compare reported allele frequencies with allele frequencies in individuals of European ancestry from the 1000 Genomes imputation reference panel^17^. We inspected P-Z plots (reported *P* value against *P* value derived from the Z score), beta and standard error distributions, and Manhattan plots to check for consistency and to identify spurious associations. For each study, variants were removed if they satisfied any one of the following criteria: imputation quality < 0.5, minor allele frequency < 0.01, absolute betas and standard errors > 10. Specific quality control measures were applied to two studies where genotyping of cases and controls was performed on different platforms, as described in Sinnott et al.^18^ and Johnson et al.^19^. To check for study-level genomic inflation, we examined quantile-quantile plots and calculated the genomic inflation factor (λ_GC_). For three studies where some degree of genomic inflation was observed (λ_GC_ > 1.1), genomic control correction was applied (**Supplementary Table 17**)^20^.

### Meta-analysis

Meta-analysis of summary data was conducted using the fixed-effect inversevariance weighted approach implemented in METAL (released March 25 2011)^21^. Variants were included if they were present in at least half of all studies. We tested for inflation of the metaanalysis test statistic due to cryptic population structure by estimating the linkage disequilibrium score regression (LDSC) intercept, implemented using LDSC v1.0.0^22^ As the LD score regression intercept indicated no inflation (LD score intercept of 1.0069), no further correction was applied to the meta-analysis summary estimates. To identify variants independently associated with HF, we analysed the genome-wide results using FUMA v1.3.2^23^, selecting a random sample of 10,000 UK Biobank participants of European ancestry as an LD reference dataset ^24^. Variants were filtered using a *P* value < 5 × 10^−8^ and independent genomic loci were LD-pruned based on an *r*^2^ < 0.1. We calculated Cochrane’s Q and *I*^2^ statistics to assess whether the effect estimates for HF sentinel variants were consistent across studies^25^.

### Heritability (*h^2^_g_*) estimation

To estimate the proportion of HF risk explained by common variants we estimated *h^2^_g_* on the liability scale using LDSC on the UK Biobank summary data (6,504 HF cases, 387,652 controls), assuming a population prevalence of 2.5%^26^. This approach assumes that a binary trait has an underlying continuous liability, and above a certain liability threshold an individual becomes affected. We can then estimate the genetic contribution to the continuous liability. Sample ascertainment can change the distribution of liability in the sampled individuals and needs to be adjusted for, which requires making assumptions about the population prevalence of the trait.

### LD reference dataset

A LD reference was created including 10,000 UKB participants of European ancestry, based on HRC-imputed genotypes (referred to henceforth as UKB10K). European individuals were identified by projecting the UK Biobank samples onto the 1000G Phase 3 samples. A genomic relationship matrix was constructed using HapMap3 variants, filtered for MAF > 0.01, pHWE < 10^−6^ and missingness < 0.05 in the European subset, and one member of each pair of samples with observed genomic relatedness greater than 0.05 was excluded to obtain a set of unrelated European individuals. Random sampling without replacement was used to extract a subset of 10,000 unrelated individuals of European ancestry. Variants with a minor allele count (MAC) > 5, a genotype probability > 0.9 and imputation quality > 0.3 were converted to hard calls. This LD reference dataset was used for downstream summary-based analysis and for identifying SNP proxies.

### Gene Mapping Analysis

#### Gene set enrichment analysis

A gene-based and gene-set enrichment analysis of variant associations was performed using MAGMA^27^, implemented by FUMA v1.3.2 ^23^. This analysis was performed using summary-level meta-analysis results. First, a gene-based association analysis to identify candidate genes associated with HF was conducted. Second, a tissue-enrichment analysis of HF associated genes was performed using gene expression data for 30 tissues from GTEx. Finally, a gene-set enrichment analysis was performed based on pathway annotations from the Gene Ontology database^28^. For all MAGMA analyses, multiple testing was accounted for by Bonferroni correction.

#### Missense consequences of sentinel variants and proxies

We queried the protein-coding consequence of the sentinel variants and proxies (r^2^ > 0.8) using the Combined Annotation Dependent Depletion (CADD) score^29^, implemented using FUMA v1.3.2^23^. The CADD score integrates information from 63 distinct functional annotations into a single quantitative score, ranging from 1 to 99, based on variant rank relative to all 8.6 billion possible SNVs of the human reference genome (GRCh37). Sentinel SNPs or proxies with CADD score > 20 were identified. A CADD score of 20 indicates that the variant is ranked in the top 1% of highest scoring variants, while a CADD score of 30 indicates the variant is ranked in the top 0.1%.

#### Expression quantitative trait analysis

To determine if HF sentinel variants had *cis*-effects on gene expression, we queried two expression quantitative trait loci (eQTL) datasets based on RNA sequencing of human heart tissue – the Genotype-Tissue Expression (GTEx) v7 resource ^30^ and the Myocardial Applied Genomics Network (MAGNet) repository (http://www.med.upenn.edu/magnet/). The GTExv7 sample included 272 left ventricular (LV) and 264 right atrium auricular (RAA) non-diseased tissue samples from European (83.7%) and African Americans (15.1%) individuals. The MAGNet repository included 89 LV and 101 LA tissue samples obtained from rejected donor tissue from hearts with no evidence of structural disease; and 89 LV samples from individuals with dilated cardiomyopathy, obtained at the time of transplantation. eQTL analysis of the LV data from MAGNet analysis was performed using the QTLtools package^31^ in DCM with adjustment for age, sex, disease status and the first 3 genetic PCs. To account for observed batch effects, a surrogate variant analysis was performed using the R package SVAseq^32^ and 22 additional covariates were identified and included in the model. eQTL datasets for LA tissue from MAGNet, and heart tissue from GTEx we derived as previously reported^33,34^. We queried HF sentinel variants for eQTL associations with genes located either fully or partly within a 1 megabase (Mb) upstream or downstream of the sentinel variant (referred to as *cis*-genes). We accounted for multiple testing by adjusting a significance threshold of *P* < 0.05 for the total number of SNP-*cis*-gene tests performed across the four heart tissue eQTL datasets (P < 4.73E-05 for a total of 1,056 SNP-gene associations). Baseline characteristics for the MAGNet study are provided in **Supplementary Table 18**. We also queried sentinel HF variants for associations with *cis*-gene expression in blood from the eQTLGen consortium (N = 31,684)^35^. Given the large sample size, we used a stringent genome-wide significance threshold of *P* < 5 × 10^−8^ to identify significant blood eQTLs.

#### Colocalisation analysis

Bayesian colocalisation analysis was performed using *coloc* to test whether shared associations with gene expression and heart failure risk were consistent with a single common causal variant hypothesis^36^. We tested all genes with significant *cis*-eQTL association by analysing all variants within a 200 kilobase window around the gene using eQTL summary data for heart tissues and whole blood, and HF summary data from present study. We set the prior probability of a SNP being associated only with gene expression, only with HF, or with both traits as 10^−4^, 10^−4^, and 10^−5^. For each gene, we report the posterior probability that the association with gene expression and HF risk is driven by a single causal variant. We consider a posterior probability of ≥ 0.7 as providing evidence supporting a causal role for the gene as a mediator of HF risk.

#### Transcriptome-wide association analysis

We employed the S-PrediXcan method^37^ implemented in the MetaXcan software (https://github.com/hakyimlab/MetaXcan) to identify genes whose predicted expression levels in heart tissue are associated with HF risk. Prediction models trained on GTExv7 heart tissue datasets were applied to the HERMES meta-analysis results. Only models that significantly predicted gene expression in the GTEx eQTL dataset (FDR <0.05) were considered. A total of 4859 genes were tested in left ventricle tissue and 4467 genes for right atrial appendage. Genes with an association P < 5.36×10-6 (0.05 / (4859 + 4467)) were considered to have gene expression profiles significantly associated with HF.

#### Protein quantitative trait analysis in blood

We queried both *cis*- and *trans*-protein QTL (pQTL) associations based on measures for serum proteins mapping to 3000 genes in 3,301 healthy individuals from the INTERVAL study^38^. We accounted for multiple testing by adjusting a significance threshold of *P* < 0.05 for the total number of tests for all variants and proteins tested (36,000 tests).

### Association of HR risk loci with other phenotypes

#### GWAS Catalog

We queried associations (with *P* < 1 × 10^−5^) of sentinel variants and proxies (r^2^ > 0.6) with any trait in the NHGRI-EBI Catalog of published GWAS (accessed 21 January 2019)^39,40^.

#### UK Biobank phenotypes

We report associations (where *P* < 1 × 10^−5^) for the sentinel variants with traits in the UK Biobank cohort using the MRBase PheWAS database (http://phewas.mrbase.org/, accessed 17 Jan 2019). The database contains GWA summary data for 4,203 phenotypes measured in 361,194 unrelated individuals of European ancestry from the UK Biobank data.

#### HF-related phenotypes

We queried GWAS data for ten traits related to HF risk factors, endophenotypes and related disease traits using summary-level data from the largest available GWAS study (either publicly available or through agreement with study investigators). The following phenotypes were considered: fractional shortening (FS), left ventricular dimension (LVD)^41^, dilated cardiomyopathy (DCM); atrial fibrillation (AF)^42^, coronary artery disease (CAD)^43^, low density lipoprotein cholesterol (LDL-C)^44^, type 2 diabetes (T2D)^45^; body mass index (BMI)^46^, systolic blood pressure (SBP), and diastolic blood pressure (DBP)^47^. For DCM, a GWAS was performed in the UKB among individuals of European ancestry with cases defined by the presence of ICD10 code I42.0 as a main/secondary diagnosis or primary/secondary cause of death with non-cases as referents, using PLINK2. Logistic regression was performed with adjustment for age, sex, genotyping array, and the first ten principal components.

#### Hierarchical agglomerative clustering

We performed hierarchical agglomerative clustering on a locus level using the complete linkage method based on the associations with related traits as described above. Where a sentinel variant is not available in any of the other traits summary results, a common proxy is used in place of the sentinel variant. For the LPA locus, we used associations for a proxy of the more common variant (rs55730499). Dissimilarity structure was calculated using Euclidean distance based on the Z-score (beta of continuous traits or log odds of disease risk divided by standard error) of the cross-trait associations. We accounted for multiple testing at family-wise error rate of 0.05 by Bonferroni correction for the 10 traits tested per HF locus (110 tests), and considered P < 4.5e^−4^ (0.05 / 110) as our significance threshold for association.

### Genetic correlation analysis

We estimated genetic correlation between HF and eleven risk factors using LD score regression^22^ on the GWAS summary statistics for each trait: AF^42^, CAD^43^, LDL-C, high density lipoprotein cholesterol (HDL-C), triglycerides (TG)^44^, T2D^45^; BMI^46^, SBP, DBP^47^, heart rate (HR)^48^ and estimated glomerular filtration rate (eGFR)^49^.

### Mendelian randomisation analysis

We performed two sample Mendelian randomisation analysis using the Generalized summary data-based Mendelian randomisation (GSMR)^50^ implemented in GCTA v1.91.7beta^51^. To identify independent SNP instruments for each exposure, GWAS-significant SNPs (*P* < 5×10^−08^) for each risk factor were pruned (*r*^2^ < 0.05; LD window of 10,000kb; using the UKB10K LD reference). We then estimated the causal effect of the risk factor on the disease trait according to the MR paradigm. The HEIDI test implemented in GSMR was used to detect and remove (if HEIDI p-value <0.01) variants showing horizontal pleiotropy i.e. having independent effects on both exposure and outcome, as such variants do not satisfy the underlying assumptions for valid instruments. As sensitivity analyses, we estimated the causal effects of known risk factors on HF risk other statistical methodology and software – the R package TwoSampleMR^52^ was used to select independent variant instruments for the exposure using the same parameters as per the GSMR analysis (*P* < 5×10^−8^; *r2* <0.05; LD window of 10,000kb), except the TwoSampleMR package uses the 1000 Genomes as the LD reference. Causal estimates based on the inverse-variance weighted (IVW)^53^, MR-Egger and median weighted methods^54^ were then calculated using the MendelianRandomization^55^ R package. To enable comparison of MR estimates between traits, we present effect estimates corresponding to the risk of HF for a 1-standard deviation (SD) higher risk factor of interest. Where the original GWAS conducted rank based inverse normal transformation (RINT) of a trait prior to GWAS, we used the per-allele beta coefficients following RINT to approximate the equivalent values on the standardised scale, as has been conducted previously.

To determine if the causal effects of the continuous risk factors on HF were mediated via their effects on CAD or AF risk, we repeated the GSMR analysis after conditioning the HF summary statistics on CAD and AF GWAS summary statistics, as described below.

#### Conditional analysis

To estimate the effects of HF risk variants after adjusting for risk factors which showed a significant causal effect on HF in the MR analyses, we performed the Multi-trait Conditional and Joint Analysis (mtCOJO) on summary data, as implemented in GCTA v1.91.7beta^51^. HF summary statistics were adjusted for AF^34^, CAD^43^, LDL-C, HDL-C, triglycerides^44^, DBP, SBP^47^ and BMI^46^ using GWAS summary data. The UKB10K LD reference was used.

#### Reporting summary

Further information is provided in the Nature Research Reporting Summary.

#### Data availability

The data sets generated during this study are available from the corresponding author upon reasonable request. The summary GWAS estimates for this analysis are available on the Cardiovascular Disease Knowledge Portal (http://www.broadcvdi.org/).

## Notes

**Competing interests** J.B.W., L.B., Xing Chen, C.L.H., M.W.N. and A. Malarstig are current or former employee of Pfizer who may hold Pfizer stock and/or stock options. J.D.B. and J.C. are employees of Regeneron Genetics Center. M.E.D. is an employee of Regeneron Pharmaceuticals. W.M. reports grants and personal fees from Siemens Diagnostics, grants and personal fees from Aegerion Pharmaceuticals, grants and personal fees from AMGEN, grants and personal fees from Astrazeneca, grants and personal fees from Danone Research, personal fees from Hoffmann LaRoche, personal fees from MSD, grants and personal fees from Pfizer, personal fees from Sanofi, personal fees from Synageva, grants and personal fees from BASF, grants from Abbott Diagnostics, grants and personal fees from Numares AG, grants and personal fees from Berlin-Chemie, employment with Synlab Holding Deutschland GmbH, all outside the submitted work. M.L.O. reports grant support from GlaxoSmithKline, Eisai, Janssen, Merck, and AstraZeneca. B.M.P. serves on the DSMB of a clinical trial funded by Zoll LifeCor and on the Steering Committee of the Yale Open Data Access Project funded by Johnson & Johnson. V.S. participated in a conference trip sponsored by Novo Nordisk and received a honorarium from the same source for participating in an advisory board meeting. He also has ongoing research collaboration with Bayer Ltd. B.T. is a full-time employee of Servier. S.A.L. receives sponsored research support from Bristol Myers Squibb / Pfizer, Bayer AG, and Boehringer Ingelheim, and has consulted for Abbott, Quest Diagnostics, Bristol Myers Squibb / Pfizer. M.V.H. has collaborated with Boehringer Ingelheim in research, and in accordance with the policy of the The Clinical Trial Service Unit and Epidemiological Studies Unit (University of Oxford), did not accept any personal payment. P.T.E. receives sponsored research support from Bayer AG, and has consulted with Bayer AG, Novartis and Quest Diagnostics. D.I.S. is a full-time employee of BenevolentAI. R.T.L. has received research grants from Pfizer. The remaining authors declare no competing interest.

## References

1. Ziaeian, B. & Fonarow, G. C. Epidemiology and aetiology of heart failure. Nature Reviews Cardiology (2016). doi:10.1038/nrcardio.2016.25

2. Lindgren, M. P. et al. A Swedish Nationwide Adoption Study of the Heritability of Heart Failure. JAMA Cardiol. 3, 703–710 (2018).

3. O’Donnell, C. J., Yancy, C. W. & McNally, E. M. Is Heart Failure Inherited?: Beyond the Cardiomyopathies, Genetics Do Matter. JAMA Cardiol. 3, 710–711 (2018).

4. Tayal, U., Prasad, S. & Cook, S. A. Genetics and genomics of dilated cardiomyopathy and systolic heart failure. Genome Med. 9, 20 (2017).

5. Roger, V. L. et al. Trends in Heart Failure Incidence and Survival in a Community-Based Population. JAMA 292, 344 (2004).

6. Ponikowski, P. et al. 2016 ESC Guidelines for the diagnosis and treatment of acute and chronic heart failure. European Heart Journal (2016). doi:10.1093/eurheartj/ehw128

7. Kenchaiah, S. et al. Obesity and the Risk of Heart Failure. N. Engl. J. Med. 347, 305–313 (2002).

8. Ziaeian, B. & Fonarow, G. C. Epidemiology and aetiology of heart failure. Nature Reviews Cardiology 13, 368–378 (2016).

9. Cahill, T. J., Ashrafian, H. & Watkins, H. Genetic Cardiomyopathies Causing Heart Failure. Circ. Res. 113, 660–675 (2013).

10. Aragam, K. G. et al. Phenotypic Refinement of Heart Failure in a National Biobank Facilitates Genetic Discovery. Circulation 139, 489–501 (2019).

11. Smith, N. L. et al. Association of genome-wide variation with the risk of incident heart failure in adults of European and African ancestry: A prospective meta-analysis from the cohorts for heart and aging research in genomic epidemiology (CHARGE) consortium. Circ. Cardiovasc. Genet. 3, 256–66 (2010).

12. Meder, B. et al. A genome-wide association study identifies 6p21 as novel risk locus for dilated cardiomyopathy. Eur. Heart J. 35, 1069–77 (2014).

13. Esslinger, U. et al. Exome-wide association study reveals novel susceptibility genes to sporadic dilated cardiomyopathy. PLoS One 12, e0172995 (2017).

14. Villard, E. et al. A genome-wide association study identifies two loci associated with heart failure due to dilated cardiomyopathy. Eur. Heart J. 32, 1065–76 (2011).

15. Davey Smith, G. & Ebrahim, S. ‘Mendelian randomization’: can genetic epidemiology contribute to understanding environmental determinants of disease?*. Int. J. Epidemiol. 32, 1–22 (2003).

16. Bulik-Sullivan, B. K. et al. LD Score regression distinguishes confounding from polygenicity in genome-wide association studies. Nat. Genet. (2015). doi:10.1038/ng.3211

17. Benjamin, E. J. et al. Heart Disease and Stroke Statistics—2018 Update: A Report From the American Heart Association. Circulation 137, e67–e492 (2018).

18. Welter, D. et al. The NHGRI GWAS Catalog, a curated resource of SNP-trait associations. Nucleic Acids Res. (2014). doi:10.1093/nar/gkt1229

19. Wild, P. S. et al. Large-scale genome-wide analysis identifies genetic variants associated with cardiac structure and function. J. Clin. Invest. 127, 1798–1812 (2017).

20. Roselli, C. et al. Multi-ethnic genome-wide association study for atrial fibrillation. Nat. Genet. 50, 1225–1233 (2018).

21. Nikpay, M. et al. A comprehensive 1,000 Genomes-based genome-wide association metaanalysis of coronary artery disease. Nat. Genet. 47, 1121–1130 (2015).

22. Warren, H. R. et al. Genome-wide association analysis identifies novel blood pressure loci and offers biological insights into cardiovascular risk. Nat. Genet. 49, 403–415 (2017).

23. Locke, A. E. et al. Genetic studies of body mass index yield new insights for obesity biology. Nature 518, 197–206 (2015).

24. Eppinga, R. N. et al. Identification of genomic loci associated with resting heart rate and shared genetic predictors with all-cause mortality. Nat. Genet. 48, 1557–1563 (2016).

25. Willer, C. J. et al. Discovery and refinement of loci associated with lipid levels. Nat. Genet. 45, 1274–1283 (2013).

26. Scott, R. A. et al. An Expanded Genome-Wide Association Study of Type 2 Diabetes in Europeans. Diabetes 66, 2888–2902 (2017).

27. Santhanakrishnan, R. et al. Atrial Fibrillation Begets Heart Failure and Vice Versa: Temporal Associations and Differences in Preserved Versus Reduced Ejection Fraction. Circulation 133, 484–92 (2016).

28. Zhu, Z. et al. Causal associations between risk factors and common diseases inferred from GWAS summary data. Nat. Commun. (2018). doi:10.1038/s41467-017-02317-2

29. de Leeuw, C. A., Mooij, J. M., Heskes, T. & Posthuma, D. MAGMA: generalized gene-set analysis of GWAS data. PLoS Comput. Biol. 11, e1004219 (2015).

30. Domínguez, F. et al. Dilated Cardiomyopathy Due to BLC2-Associated Athanogene 3 (BAG3) Mutations. J. Am. Coll. Cardiol. 72, 2471–2481 (2018).

31. Võsa, U. et al. Unraveling the polygenic architecture of complex traits using blood eQTL meta-analysis. bioRxiv (2018).

32. Giambartolomei, C. et al. Bayesian Test for Colocalisation between Pairs of Genetic Association Studies Using Summary Statistics. PLoS Genet. 10, e1004383 (2014).

33. Sun, B. B. et al. Genomic atlas of the human plasma proteome. Nature 558, 73–79 (2018).

34. Frey, N. et al. Calsarcin-2 deficiency increases exercise capacity in mice through calcineurin/NFAT activation. J. Clin. Invest. 118, 3598–608 (2008).

35. Molkentin, J. D. Parsing good versus bad signaling pathways in the heart: role of calcineurin-nuclear factor of activated T-cells. Circ. Res. 113, 16–9 (2013).

36. Beqqali, A. et al. CHAP is a newly identified Z-disc protein essential for heart and skeletal muscle function. J. Cell Sci. 123, 1141–50 (2010).

37. Behl, C. Breaking BAG: The Co-Chaperone BAG3 in Health and Disease. Trends Pharmacol. Sci. 37, 672–688 (2016).

38. Tane, S. et al. CDK inhibitors, p21Cip1 and p27Kip1, participate in cell cycle exit of mammalian cardiomyocytes. Biochem. Biophys. Res. Commun. 443, 1105–1109 (2014).

39. Mattioli, E. et al. Altered modulation of lamin A/C-HDAC2 interaction and *p21* expression during oxidative stress response in HGPS. Aging Cell e12824 (2018). doi:10.1111/acel.12824

40. Boyden, L. M. et al. Mutations in kelch-like 3 and cullin 3 cause hypertension and electrolyte abnormalities. Nature 482, 98–102 (2012).

41. Sciarretta, S., Palano, F., Tocci, G., Baldini, R. & Volpe, M. Antihypertensive Treatment and Development of Heart Failure in Hypertension. Arch. Intern. Med. 171, 384–94 (2011).

42. Velagaleti, R. S. & Vasan, R. S. Heart failure in the twenty-first century: is it a coronary artery disease or hypertension problem? Cardiol. Clin. 25, 487–95; v (2007).

43. Roger, V. L. Epidemiology of Heart Failure. Circ. Res. 113, 646–659 (2013).

44. Fry, A. et al. Comparison of Sociodemographic and Health-Related Characteristics of UK Biobank Participants With Those of the General Population. Am. J. Epidemiol. 186, 1026–1034 (2017).

45. He, L. et al. Causal effects of cardiovascular risk factors on onset of major age-related diseases: A time-to-event Mendelian randomization study. Exp. Gerontol. 107, 74–86 (2018).

46. Fall, T. et al. The role of adiposity in cardiometabolic traits: a Mendelian randomization analysis. PLoS Med. 10, e1001474 (2013).

47. Dhingra, R., Gaziano, J. M. & Djoussé, L. Chronic kidney disease and the risk of heart failure in men. Circ. Heart Fail. 4, 138–44 (2011).

48. Nanchen, D. et al. Resting Heart Rate and the Risk of Heart Failure in Healthy Adults. Circ. Hear. Fail. 6, 403–410 (2013).

49. Roselli, C. et al. Multi-ethnic genome-wide association study for atrial fibrillation. Nat. Genet. 50, 1225–1233 (2018).

## Online Methods References

1. The 1000 Genomes Project Consortium. A map of human genome variation from population-scale sequencing. Nature 467, 1061–73 (2010).

2. International HapMap Consortium et al. A second generation human haplotype map of over 3.1 million SNPs. Nature 449, 851–61 (2007).

3. McCarthy, S. et al. A reference panel of 64,976 haplotypes for genotype imputation. Nat. Genet. (2016). doi:10.1038/ng.3643

4. Mitt, M. et al. Improved imputation accuracy of rare and low-frequency variants using population-specific high-coverage WGS-based imputation reference panel. Eur. J. Hum. Genet. 25, 869–876 (2017).

5. Loh, P.-R., Palamara, P. F. & Price, A. L. Fast and accurate long-range phasing in a UK Biobank cohort. Nat. Genet. 48, 811–816 (2016).

6. Li, Y., Willer, C. J., Ding, J., Scheet, P. & Abecasis, G. R. MaCH: using sequence and genotype data to estimate haplotypes and unobserved genotypes. Genet. Epidemiol. 34, 816–834 (2010).

7. Delaneau, O., Zagury, J.-F. & Marchini, J. Improved whole-chromosome phasing for disease and population genetic studies. Nat. Methods 10, 5–6 (2013).

8. Fuchsberger, C., Abecasis, G. R. & Hinds, D. A. minimac2: faster genotype imputation. Bioinformatics 31, 782–784 (2015).

9. Howie, B. N., Donnelly, P. & Marchini, J. A Flexible and Accurate Genotype Imputation Method for the Next Generation of Genome-Wide Association Studies. PLoS Genet. 5, e1000529 (2009).

10. Kong, A. et al. Detection of sharing by descent, long-range phasing and haplotype imputation. Nat. Genet. 40, 1068–1075 (2008).

11. Aulchenko, Y. S., Struchalin, M. V & van Duijn, C. M. ProbABEL package for genome-wide association analysis of imputed data. BMC Bioinformatics 11, 134 (2010).

12. Kutalik, Z. et al. Methods for testing association between uncertain genotypes and quantitative traits. Biostatistics 12, 1–17 (2011).

13. Chang, C. C. et al. Second-generation PLINK: Rising to the challenge of larger and richer datasets. Gigascience (2015). doi:10.1186/s13742-015-0047-8

14. Marchini, J., Howie, B., Myers, S., McVean, G. & Donnelly, P. A new multipoint method for genome-wide association studies by imputation of genotypes. Nat. Genet. (2007). doi:10.1038/ng2088

15. R Core team. R Core Team. R: A Language and Environment for Statistical Computing. R Foundation for Statistical Computing, Vienna, Austria. ISBN 3-900051-07-0, URL http://www.R-project.org/. (2015).

16. Winkler, T. W. et al. Quality control and conduct of genome-wide association meta-analyses. Nat. Protoc. (2014). doi:10.1038/nprot.2014.071

17. Auton, A. et al. A global reference for human genetic variation. Nature (2015). doi:10.1038/nature15393

18. Sinnott, J. A. & Kraft, P. Artifact due to differential error when cases and controls are imputed from different platforms. Hum. Genet. 131, 111–119 (2012).

19. Johnson, E. O. et al. Imputation across genotyping arrays for genome-wide association studies: assessment of bias and a correction strategy. Hum. Genet. 132, 509–522 (2013).

20. Devlin, B. & Roeder, K. Genomic control for association studies. Biometrics 55, 997–1004 (1999).

21. Willer, C. J., Li, Y. & Abecasis, G. R. METAL: Fast and efficient meta-analysis of genomewide association scans. Bioinformatics (2010). doi:10.1093/bioinformatics/btq340

22. Bulik-Sullivan, B. K. et al. LD Score regression distinguishes confounding from polygenicity in genome-wide association studies. Nat. Genet. (2015). doi:10.1038/ng.3211

23. Watanabe, K., Taskesen, E., Van Bochoven, A. & Posthuma, D. Functional mapping and annotation of genetic associations with FUMA. Nat. Commun. (2017). doi:10.1038/s41467-017-01261-5

24. Bycroft, C. et al. The UK Biobank resource with deep phenotyping and genomic data. Nature 562, 203–209 (2018).

25. Higgins, J. P. T., Thompson, S. G., Deeks, J. J. & Altman, D. G. Measuring inconsistency in meta-analyses. BMJ Br. Med. J. (2003). doi:10.1136/bmj.327.7414.557

26. Benjamin, E. J. et al. Heart Disease and Stroke Statistics—2018 Update: A Report From the American Heart Association. Circulation 137, e67–e492 (2018).

27. de Leeuw, C. A., Mooij, J. M., Heskes, T. & Posthuma, D. MAGMA: generalized gene-set analysis of GWAS data. PLoS Comput. Biol. 11, e1004219 (2015).

28. Ashburner, M. et al. Gene Ontology: tool for the unification of biology. Nat. Genet. 25, 25–29 (2000).

29. Kircher, M. et al. A general framework for estimating the relative pathogenicity of human genetic variants. Nat. Genet. 46, 310–5 (2014).

30. GTEx Consortium. The Genotype-Tissue Expression (GTEx) project. Nat. Genet. 45, 580–5 (2013).

31. Delaneau, O. et al. A complete tool set for molecular QTL discovery and analysis. Nat. Commun. 8, 15452 (2017).

32. Leek, J. T. svaseq: removing batch effects and other unwanted noise from sequencing data. Nucleic Acids Res. 42, (2014).

33. GTEx Consortium et al. Genetic effects on gene expression across human tissues. Nature 550, 204–213 (2017).

34. Roselli, C. et al. Multi-ethnic genome-wide association study for atrial fibrillation. Nat. Genet. 50, 1225–1233 (2018).

35. Võsa, U. et al. Unraveling the polygenic architecture of complex traits using blood eQTL meta-analysis. bioRxiv (2018).

36. Giambartolomei, C. et al. Bayesian Test for Colocalisation between Pairs of Genetic Association Studies Using Summary Statistics. PLoS Genet. 10, e1004383 (2014).

37. Barbeira, A. N. et al. Exploring the phenotypic consequences of tissue specific gene expression variation inferred from GWAS summary statistics. Nat. Commun. 9, 1825 (2018).

38. Sun, B. B. et al. Genomic atlas of the human plasma proteome. Nature 558, 73–79 (2018).

39. Welter, D. et al. The NHGRI GWAS Catalog, a curated resource of SNP-trait associations. Nucleic Acids Res. (2014). doi:10.1093/nar/gkt1229

40. MacArthur, J. et al. The new NHGRI-EBI Catalog of published genome-wide association studies (GWAS Catalog). Nucleic Acids Res. (2017). doi:10.1093/nar/gkw1133

41. Wild, P. S. et al. Large-scale genome-wide analysis identifies genetic variants associated with cardiac structure and function. J. Clin. Invest. 127, 1798–1812 (2017).

42. Roselli, C. et al. Multi-ethnic genome-wide association study for atrial fibrillation. Nat. Genet. 50, 1225–1233 (2018).

43. Nikpay, M. et al. A comprehensive 1,000 Genomes-based genome-wide association metaanalysis of coronary artery disease. Nat. Genet. 47, 1121–1130 (2015).

44. Willer, C. J. et al. Discovery and refinement of loci associated with lipid levels. Nat. Genet. 45, 1274–1283 (2013).

45. Scott, R. A. et al. An Expanded Genome-Wide Association Study of Type 2 Diabetes in Europeans. Diabetes 66, 2888–2902 (2017).

46. Locke, A. E. et al. Genetic studies of body mass index yield new insights for obesity biology. Nature 518, 197–206 (2015).

47. Warren, H. R. et al. Genome-wide association analysis identifies novel blood pressure loci and offers biological insights into cardiovascular risk. Nat. Genet. 49, 403–415 (2017).

48. Eppinga, R. N. et al. Identification of genomic loci associated with resting heart rate and shared genetic predictors with all-cause mortality. Nat. Genet. 48, 1557–1563 (2016).

49. Gorski, M. et al. 1000 Genomes-based meta-analysis identifies 10 novel loci for kidney function. Sci. Rep. 7, 45040 (2017).

50. Zhu, Z. et al. Causal associations between risk factors and common diseases inferred from GWAS summary data. Nat. Commun. (2018). doi:10.1038/s41467-017-02317-2

51. Yang, J., Lee, S. H., Goddard, M. E. & Visscher, P. M. GCTA: a tool for genome-wide complex trait analysis. Am. J. Hum. Genet. 88, 76–82 (2011).

52. Hemani, G. et al. The MR-Base platform supports systematic causal inference across the human phenome. Elife (2018). doi:10.7554/eLife.34408

53. Burgess, S., Butterworth, A. & Thompson, S. G. Mendelian randomization analysis with multiple genetic variants using summarized data. Genet. Epidemiol. (2013). doi:10.1002/gepi.21758

54. Bowden, J., Davey Smith, G. & Burgess, S. Mendelian randomization with invalid instruments: effect estimation and bias detection through Egger regression. Int. J. Epidemiol. 44, 512–525 (2015).

55. Yavorska, O. O. & Burgess, S. MendelianRandomization: An R package for performing Mendelian randomization analyses using summarized data. Int. J. Epidemiol. (2017). doi:10.1093/ije/dyx034

