## Supplementary Note for "Genome-wide association study provides new insights into the genetic architecture and pathogenesis of heart failure"

### Supplementary Figures


1. Quantile-quantile plot of meta-analysis
2. Regional association plots
3. Forest plots showing association between sentinel variants and heart failure risk across study samples
4. HF risk variant effects conditioned for atrial fibrillation, coronary artery disease and blood pressure
5. Tissue enrichment analysis based on gene expression
6. Mendelian randomisation analyses for cardiovascular risk traits and HF risk
7. Mendelian randomisation funnel plots for cardiovascular risk traits and HF risk


### Supplementary Note


1. Description of candidate genes at HF risk loci
2. Description of participating studies
3. Description of participating consortia
4. Acknowledgements


### Supplementary references

### Supplementary Figures

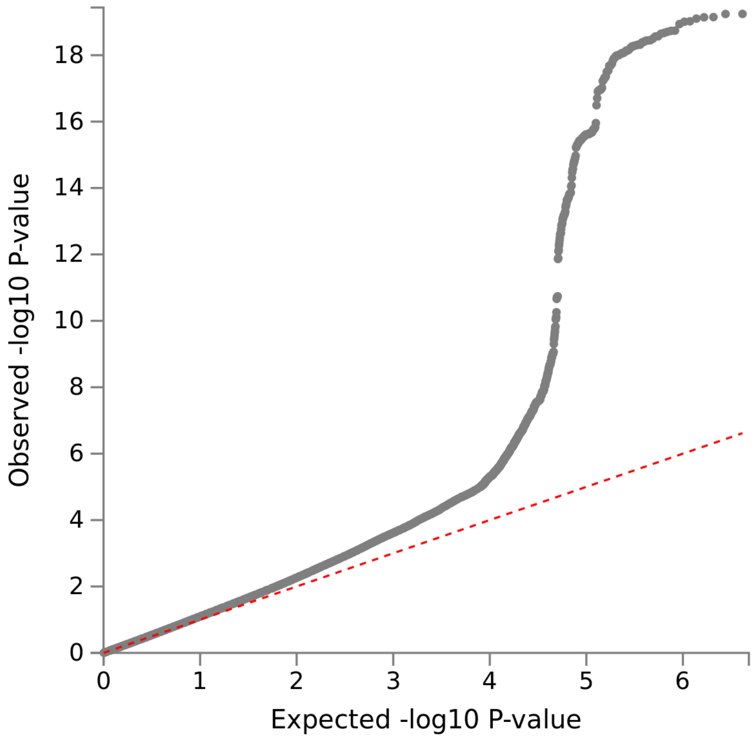


**Supplementary Figure 1**. **Quantile-quantile plot of meta-analysis.** Quantile-quantile plot of the association *P* values for 8,281,262 variants.

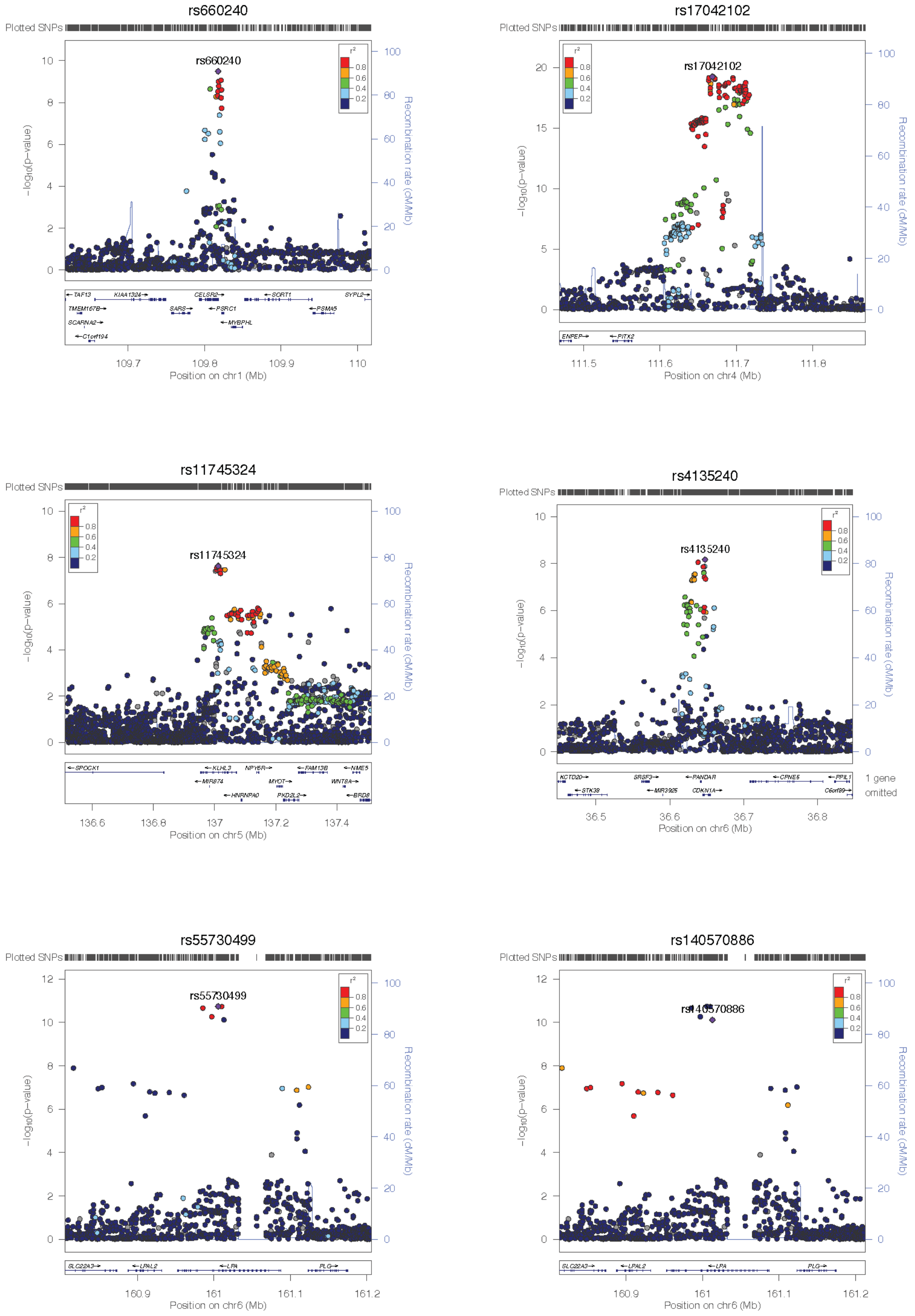


**Supplementary Figure 2. Regional association plots.** Regional association plots for each of the 12 independently associated loci in the Stage 1 meta-analysis. The x-axis represents a 1Mb region, 500kb either side of the sentinel variant (purple diamonds) and the y-axis shows –log$_{10}$ *P* values for individual SNPs. Pairwise LD ($r^2$) with the sentinel variant is based on 1000 Genomes phase 3 v5 European reference samples and is described using the colour scale given. The bottom panel show genes located within the region.

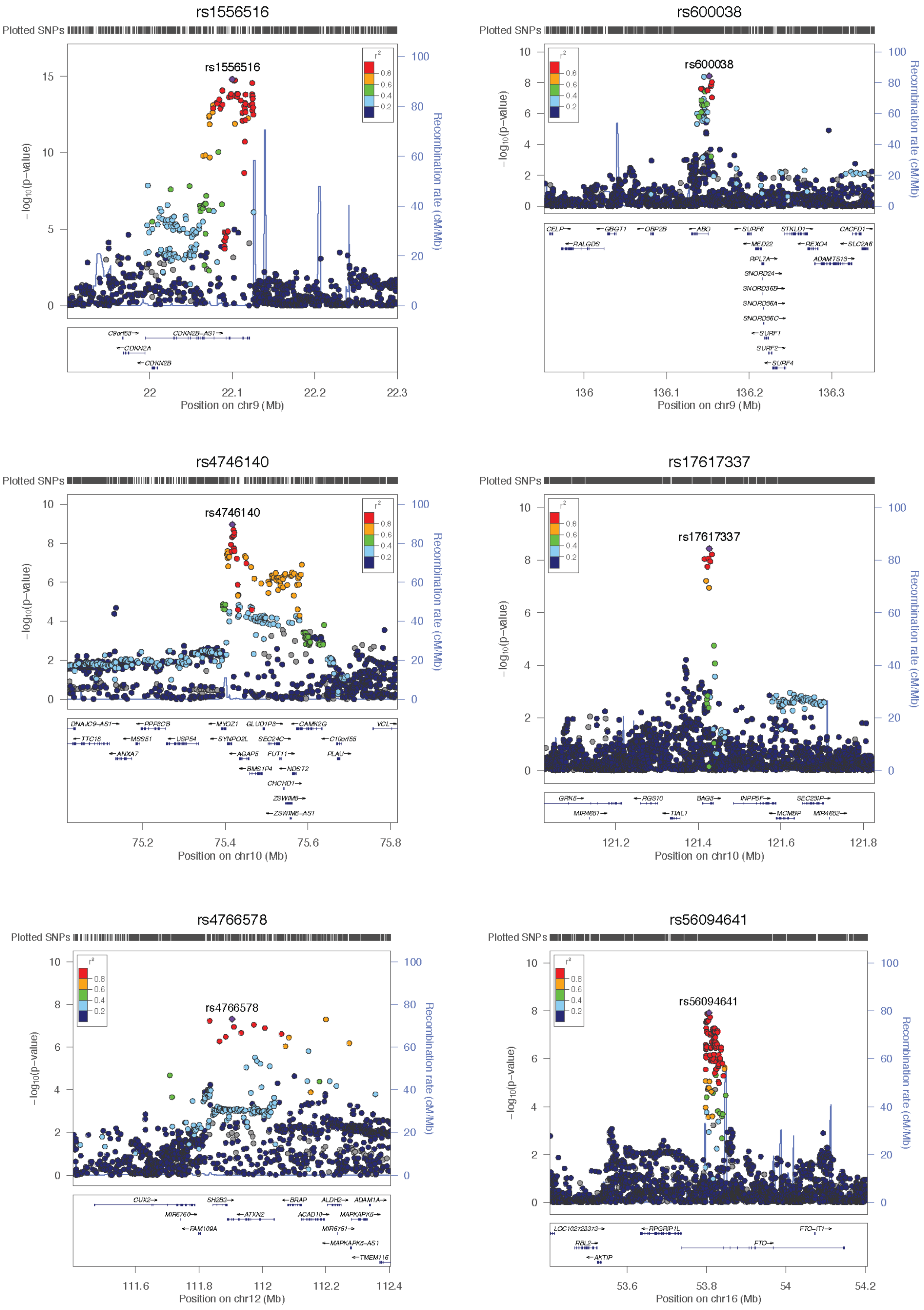


**Supplementary Figure 2. Regional association plots (cont.)**

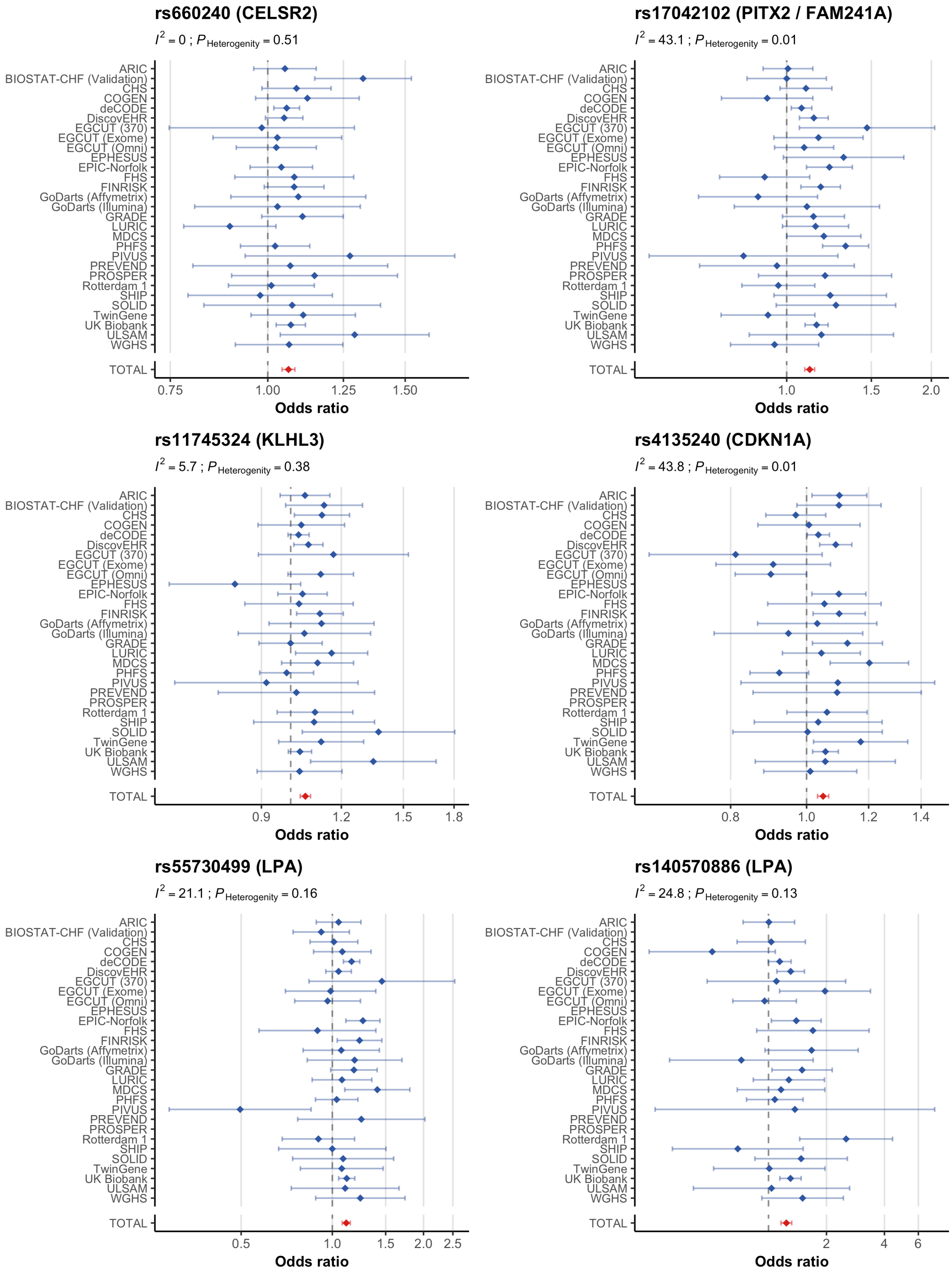


**Supplementary Figure 3. Forest plots showing association between sentinel variants and heart failure risk across study samples.** Blue squares represent the point estimate of the odds ratio and have areas proportional to the study size. The red diamond represents the point estimate for the odds ratio for the combined meta-analysis. 95% confidence intervals are shown by the width of the blue lines (individual samples) or width of the diamond (overall meta-analysis). These show the effect of each of the 12 sentinel variants on HF risk across the different studies. The I² and heterogeneity *P* values are given for each variant.

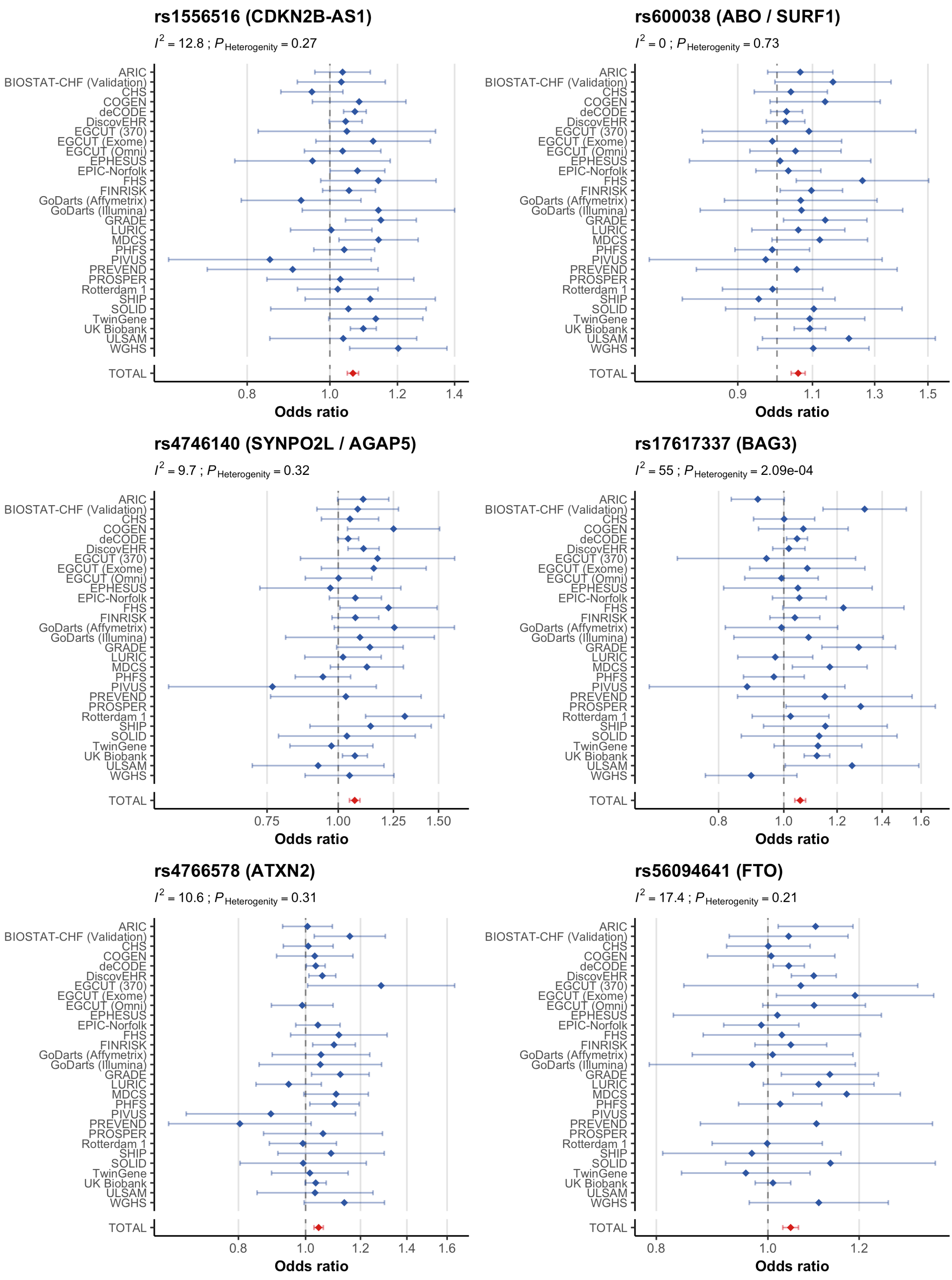


**Supplementary Figure 3. Forest plots showing association between sentinel variants and heart failure risk across study samples (cont.)**

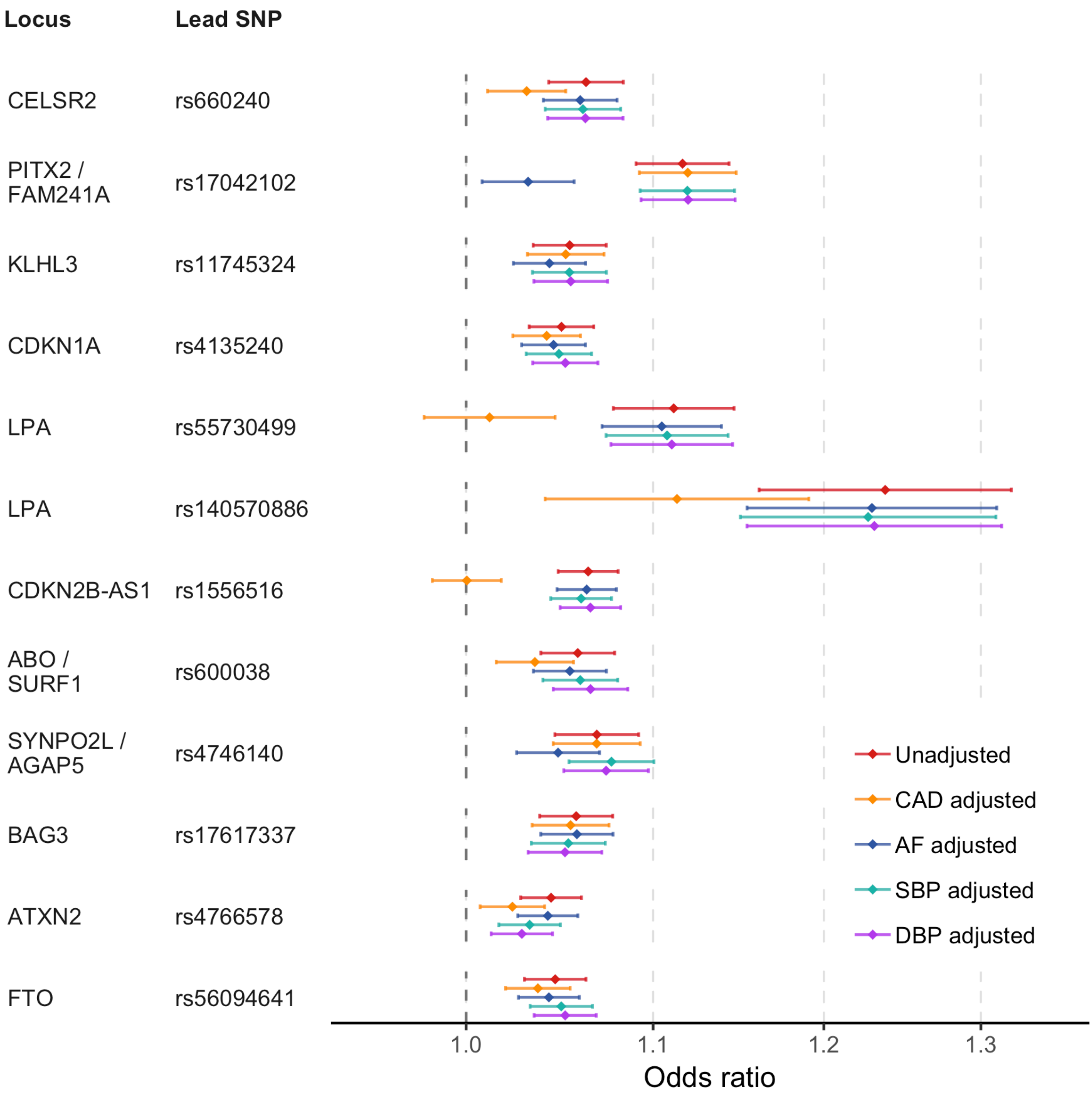


**Supplementary Figure 4. HF risk variant effects conditioned for atrial fibrillation, coronary artery disease and blood pressure.** HF sentinel variant effects are given before and after conditioning on atrial fibrillation, coronary artery disease, systolic blood pressure, and diastolic blood pressure. Conditional analysis was performed on the Stage 1 meta-analysis results and the published GWAS summary data, using GCTA-mtCOJO. AF, atrial fibrillation; CAD, coronary artery disease; SBP, systolic blood pressure; DBP, diastolic blood pressure.

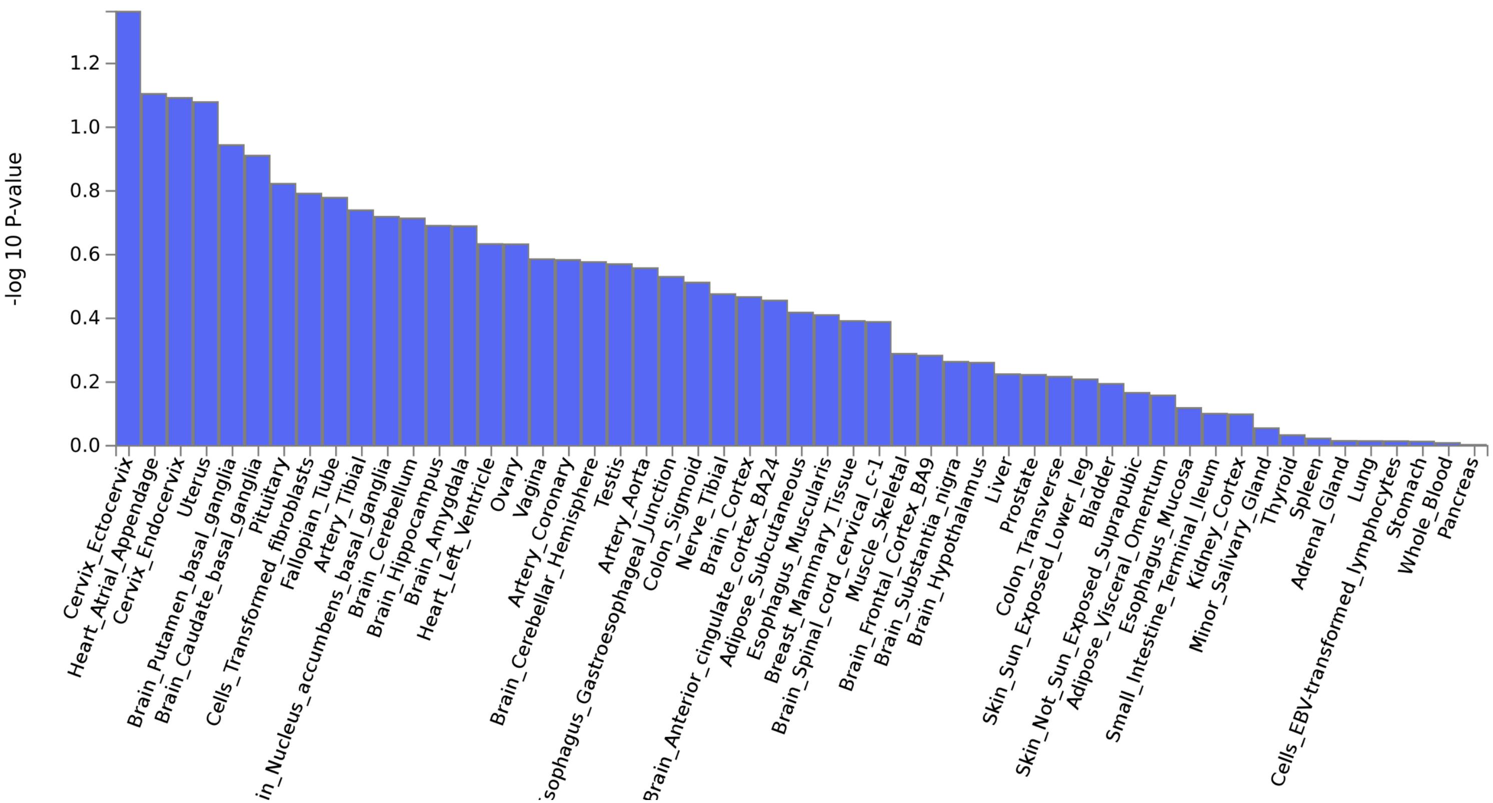


**Supplementary Figure 5. Tissue enrichment analysis based on gene expression.** Tissue enrichment analysis was performed with MAGMA, using gene expression data for 30 tissues from GTEx and gene-based association statistics based on the heart failure genome-wide meta-analysis. The y-axis shows $-\log_{10}$ *P* values for enrichment for each tissue

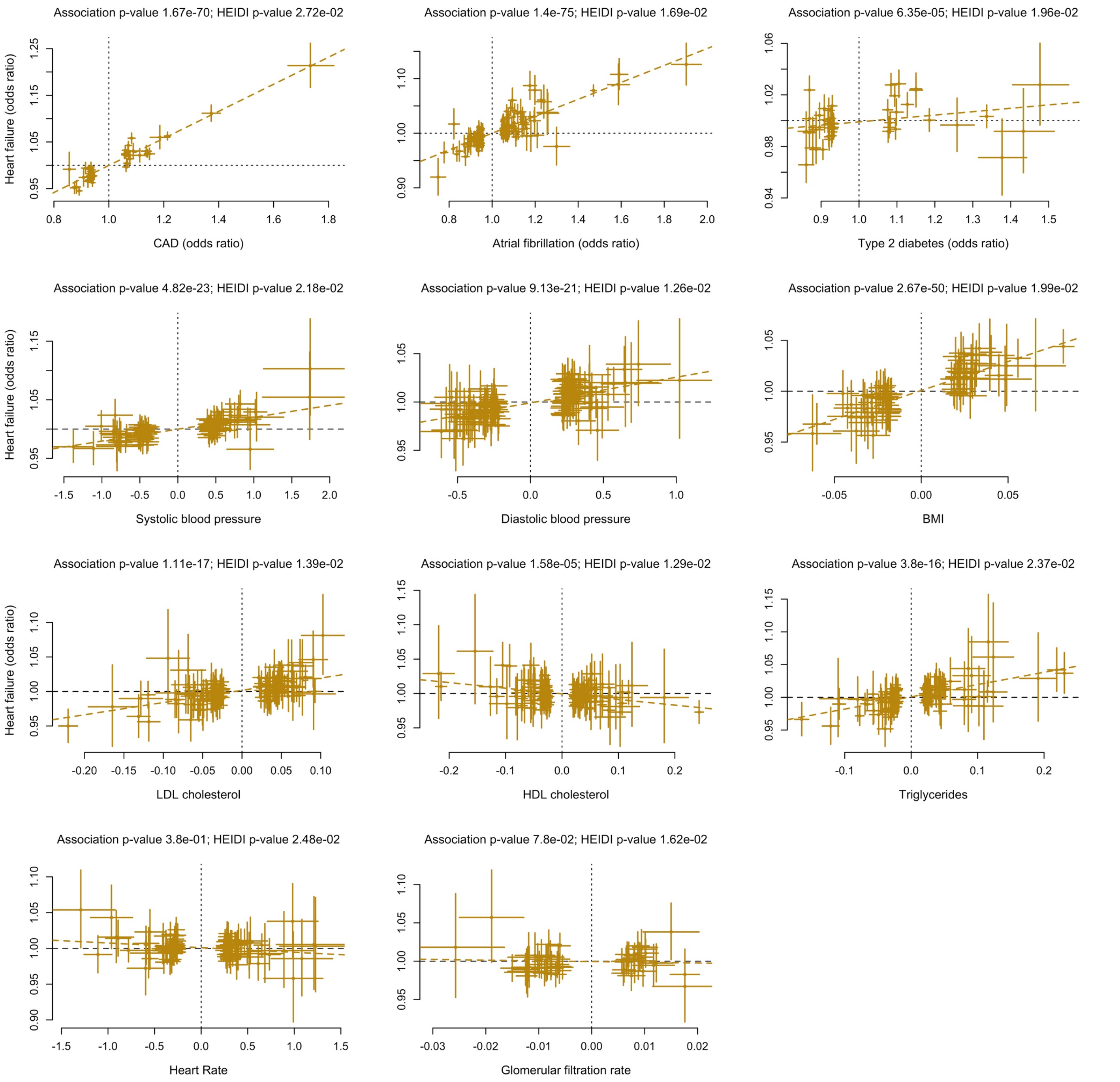


**Supplementary Figure 6. Mendelian randomisation analyses for cardiovascular risk traits and HF risk.** The plots show the effect sizes and standard errors (error bars) for the associations of independent SNP instruments for the exposure of interest and heart failure. SNP effects on the exposure($b_{zx}$) are shown on the x-axis, while the effects on HF risk are shown on the y-axis ($b_{zy}$). Units for $b_{zx}$ for blood pressure traits, heart rate and glomerular filtration rate are mmHg, beats per minute and log (ml/min/1.73 m2), respectively, while units for BMI (body mass index) and lipids are per standard deviation. Taking into account multiple-testing in 8 traits, $P < 6.25 \times 10^{-3}$ was considered as a significant causal effect of exposure on HF. Abbreviations: CAD, coronary artery disease; LDL, low density lipoprotein; HDL, high density lipoprotein.

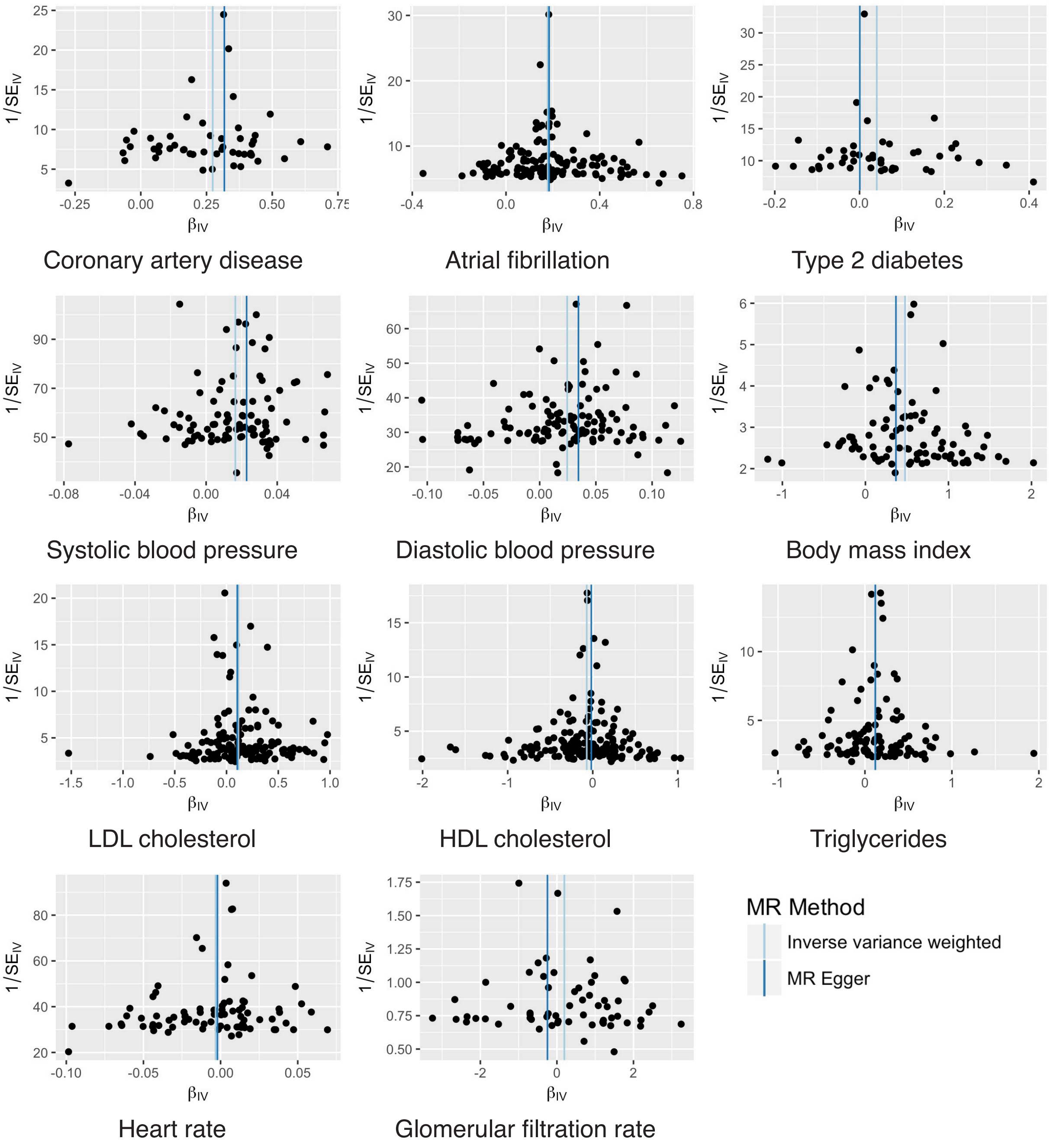


**Supplementary Figure 7. Mendelian randomisation funnel plots for cardiovascular risk traits and HF risk.** The plot shows Mendelian randomisation (MR) effect estimates ($\beta_{IV}$) per single nucleotide polymorphism (SNP) for the association between each risk trait and heart failure. The SNP $\beta_{IV}$ are plotted on the x-axis and strength of the SNP instrument, measured as the inverse of standard error unit ($1/SE_{IV}$), is given on the y-axis. Abbreviations: LDL, low density lipoprotein; HDL, high density lipoprotein.

### Supplementary Note

#### 1. Description of candidate genes at HF risk loci

***CELSR2* (rs660240)**

The sentinel variant at this locus is located in the 3' untranslated region of *CELSR2* which encodes Cadherin EGF LAG Seven-Pass G-Type Receptor 2 (CELSR2), a member of the cadherin superfamily. Variants in high LD with the sentinel variant ($r^2 \geq 0.8$) that span the *CELSR2-PSRC1-SORT1* gene cluster have been associated with cardiovascular risk factors such as LDL[1] and outcomes such as coronary artery disease[2,3] and myocardial infarction (**Supplementary Table 3**). We performed mtCOJO to estimate the effect of this locus on HF risk conditioned for LDL and CAD (**Supplementary Table 5 and Supplementary Figure 4**) and observed some attenuation of the association, though the signal was not ablated, suggesting there could be an additional effect on HF risk independent of LDL-C and CAD at this locus. eQTL analysis of blood and heart tissue revealed association of the sentinel variant with *cis*-expression of *PSRC1* but not *CELSR2* or *SORT1* in LV and blood. The posterior probability of colocalization was 0.97 in both tissues implicating this as a candidate gene for this locus (**Supplementary Tables 8 and 9**). *PSRC1* binds microtubules and is required for chromosomal segregation during mitosis[4]. This locus has been subject to extensive functional analysis and both *PSRC1* and also *SORT1* have been implicated as possible causal genes[5,6].

***PITX2 / FAM24A1* (rs17042102)**

The sentinel variant in this region, rs17042102, is located in an intergenic region between the transcription factor encoding gene, *PITX2* (paired like homeodomain 2) and *FAM241A* (encodes an uncharacterized protein). At least four independent variants at this locus are reported to be associated with AF[7]. We performed a conditional analysis for AF and demonstrated almost complete attenuation of the HF risk association (**Figure 3**). Functional studies have implicated *PITX2c,* the dominant isoform in the left atrium, at this locus. *PITX2* has a role in heart looping during development and downregulation predisposes to AF[7], and has shown to promote cardiac repair[8].

#### *KLHL3* (rs11745324)

The sentinel variant at this locus is located within the intronic region of *KLHL3* and is associated with expression of this gene in blood but not heart tissue. The HF risk allele at rs11745324 was associated with lower expression of KLHL3 in blood (**Supplementary Table 9**). The encoded protein, kelch like family member 3, is a substrate specific adaptor of the KLHL3-CUL3 (Cullin-3) ubiquitin ligase complex that negatively regulates the activity of the $Na^{+}Cl^{-}$ cotransporter (NCC; approved symbol SLC12A3) in the distal nephron through ubiquitination and proteasomal or selective autophagic degradation of NNC activators, WNK kinase 1 (WNK1) and 4 (WNK4)[9,10]. Loss of function variants in KLHL3, and other components of the pathway regulating NCC, are a cause of Familial Hyperkalaemic Hypertension (FHHt), also known as pseudohypoaldosteronism type 2 or Gordon's syndrome[11,12]. A constitutive increase in sodium and chloride reabsorption in the distal nephron, regardless of volume status, cause hypertension and reduce luminal electronegativity, leading to reduced potassium secretion. Treatment with thiazide diuretics, pharmacological inhibitors of NCC, are effective at correcting the electrolyte abnormalities and treating hypertension in FHHt. The sentinel *KLHL3* variant is also associated with AF, however the HF risk association was robust to conditioning on this related trait (**Supplementary Figure 4**).

#### *CDKN1A* (rs4135240)

The lead variant at this locus, rs4135240, is located within an intron of *CDKN1A* which encodes the cyclin-dependent kinase inhibitor p21. The minor allele at rs4135240 was associated with a lower gene expression in whole blood and a decreased risk of HF. We estimate the posterior probability of a common causal variant underlying gene expression in blood and HF as 0.71 providing evidence for a causal role of this gene. p21 is a potent cell cycle inhibitor, transcriptionally activated by p53 (TP53) in response to cell stress and DNA damage, that promotes G1 cell cycle arrest and cellular senescence[13,14]. In mice, p21 is upregulated in the first post-natal week and has been implicated in cardiomyocyte cell-cycle arrest[15]. Genetic deletion of p21 is reported to enhance regeneration of injured neonatal hearts[15,16] and of adult appendage tissue, such as ear hole puncture wounds[17]. A variant in high LD (rs733590, $r^2 = 0.91$) with the sentinel variant was associated with fractional shortening (a measure of left ventricular systolic function; $P = 1.66 \times 10^{-05}$) and AF ($P = 0.0026$), (**Supplementary Table 4**). p21 is implicated as a mediator of dysregulated stress responses and senescence in Hutchinson–Gilford progeria, a severe LMNA-linked syndrome associated with cardiovascular disease and premature aging[18,19]. Conditional analysis showed that the association of rs4135240 with HF was not explained by associations with AF.

***LPA* (rs55730499, rs140570886)**

Two variants in the intronic region of *LPA,* rs55730499 and rs140570886 were independently associated with HF and were in high linkage disequilibrium with previously reported independent signals at this locus[20]. A previous genome-wide haplotype association study identified this region as a risk locus for CAD[21]. However, the SNPs investigated in this study (rs7767084 and rs10755578) are not in LD with the HF lead SNPs ($R^2$ in Europeans < 0.05). *LPA* encodes apolipoprotein(a) which covalently binds LDL to form lipoprotein(a) (Lp(a)). Plasma Lp(a) concentration is under strong genetic control, with heritability estimates from twin studies exceeding 90%[22]. A Mendelian randomisation study using multiple independent genetic variants around the *LPA* gene as instruments for plasma Lp(a) levels, showed higher levels may play a causal role in CAD[23]. We observed complete attenuation of the association with HF for rs55730499, and only partial attenuation for rs140570886, after conditioning on LDL-C and CAD (**Supplementary Table 5 and Supplementary Figure 4**). Interestingly, rs140570886 alone was associated with expression of *SLC22A1* in blood (**Supplementary Table 9**), a gene encoding the organic cation transporter 1

(OCT1) which is expressed in cardiomyocytes and transports hormones (such as adrenaline, noradrenaline and dopamine) and drugs, therefore influencing drug disposition and response[24].

***CDKN2B-AS1* (rs7859727)**

The sentinel variant at this locus, rs7859727, is located within the intronic region of the anti-sense RNA, *CDKN2B-AS1*, also known as *ANRIL*. This gene is within the CDKN2B-CDKN2A gene cluster at chromosome 9p21, an important susceptibility locus for cardiovascular disease[9,10]. Two haplotype blocks at this locus have been shown to pre-dispose either to atherosclerosis or to myocardial infarction among individuals with pre-existing CAD[25]P is in high LD with the former block ($r^2$ 0.86 with tagging SNP rs1333049), but not with the latter block ($r^2$ 0.26 with tagging SNP rs518394). This locus has been the focus of intense study, however the causal genes mediating the risk effects at this locus remain uncertain. The expression of *CDKN2B-AS1* has been linked to variants at this locus and it may be involved in the post-transcriptional silencing of other genes at the locus, including *CDKN2B*[26]. The HF association of the sentinel risk variant at 9p21 was completely attenuated upon conditioning for CAD, suggest that this risk trait entirely mediates HF-risk at this locus.

***ABO / SURF1* (rs600038)**

The sentinel variant at this locus is located in an intergenic region between *ABO* and *SURF1*. The lead variant in this region is associated with *ABO* gene expression in blood and heart tissues, as well as ABO serum protein concentration. Colocalisation analysis implicated *ABO* as the likely causal gene at this locus in blood and heart tissues (posterior probability LA 0.83, blood 0.96. Alleles of *ABO* encode alpha 1-3-N-acetylgalactosaminyltransferase (transferase A), alpha 1-3-galactosyltransferase (transferase B), or the O allele that lacks glycosyltransferase activity due to a frameshift mutation, that determines blood group assignment. Since cardiovascular risk is associated with *ABO* blood group, we sought to determine whether the sentinel HF-risk variant at this locus was in linkage disequilibrium with variants that determine *ABO* blood group types. rs600038 (MAF 0.21) was correlated with rs8176719 (MAF 0.39, D' 0.99, $r^2$ 0.41), an indel where individuals who are homozygous for the deletion have blood group type O, rs56392308 (MAF 0.094, D' 0.9515, $r^2$ 0.0261), which determines blood group type A2 and rs8176746 (MAF 0.0845, D' 1, $r^2$ 0.025) which tags blood group type B[27]. The deletion at rs8176719 is correlated with the T allele at rs600038 which is associated with reduced HF risk, consistent with studies suggesting a reduction in cardiovascular risk with O blood types[27]. The *ABO* locus is notably pleiotropic and recent GWAS discoveries have highlighted associations with several transcripts, serum proteins and disease outcomes such as myocardial infarction[28] (**Supplementary Tables 2-3, 9, and 11**).

***SYNPO2L / AGAP5*** **(rs4746140)**

The sentinel variant is found in the intronic region of an uncharacterised non-coding RNA (ncRNA), RP11-464F9.21 which is located between two genes *SYNPO2L* (upstream) and *AGAP5* (downstream). *SYNPO2L* has an essential role in striated muscle development and function. SNPs associated with HF were also associated with expression of *MYOZ1* in heart tissue, a gene located upstream of *SYNPO2L.* Both *SYNPO2L* and *MYOZ1* are Z-disc associated cytoskeletal proteins, expressed in cardiac and skeletal muscle with a role in sarcomere organisation[29–31]. *MYOZ1* encodes myozenin 1 (also known as calsarcin-2) and is known to interact with α-actinin, filamin C, calcineurin and telethonin[32,33]. *MYOZ1* inhibits calcineurin signalling and muscle regeneration, and influences the ratio of fast and slow twitch fibres in skeletal muscle[31]. *SYNPO2L* encodes synaptopodin 2 -like (also known as cytoskeletal heart-enriched actin-associated protein, CHAP), which binds α-actinin-2, at the Z-disc, and positively regulates stress fiber assembly; its genetic deletion in zebrafish leads to reduced cardiac contractility[29,30]. eQTL analysis showed that both of these genes were expressed in atrial heart tissues. We found strong evidence of for colocalisation for *MYOZ1* in left atrium (posterior probability = 0.91) and moderate evidence for *SYNPO2L* in atrial appendage (posterior probability = 0.61) and we were unable to exclude the possibility that both are co-regulated at this locus (**Supplementary Table 8**). Of note, the syntenic organization of the close paralogues of these genes is conserved in humans and across species. For example, *SYNPO2* and *MYOZ2* on chromosome 4 and *SYNPO* and *MYOZ3* on chromosome 5. In a small candidate gene study of dilated cardiomyopathy *MYOZ1* was not implicated in this disease[34], however a rare variant in the paralog *MYOZ2* may be a causative for hypertrophic cardiomyopathy[35].

#### *BAG3* (rs17617337)

The sentinel variant at this locus is located in the intronic region of the *BAG3* gene which encodes BCL2 associated athanogene 3 (BAG3). Previous reports indicate that BAG3 is highly expressed in cardiac and skeletal muscle and colocalises with α-actinin and desmin at Z discs[36]. The sentinel HF-risk variant at this locus, rs17617337, was not associated with *BAG3* expression in heart tissues however, we found that the HF-risk allele was associated with decreased gene expression and protein abundance in whole blood (**Supplementary Tables 9 and 11**). A large number of non-synonymous *BAG3* variants have been associated with dilated cardiomyopathy (DCM)[37–40], and early onset severe axonal neuropathy[41]. Furthermore, the common non-synonymous variant rs2234962 identified in a GWAS of DCM is in high LD with the HF-risk allele reported here ($r^2$ = 0.99)[42]. Decreased levels of the BAG3 protein have also been reported in dilated cardiomyopathy and heart failure[43] and increased expression of BAG3 is induced by heat shock factor 1 in response to cellular stress[44]. Mice with a homozygous *Bag3* deletion ($Bag3^{-/-}$) developed non-inflammatory myofibrillar degeneration, myocyte apoptosis and cardiomyopathy[36] and haploinsufficiency ($Bag3^{+/-}$) was associated with increased apoptosis and progressive left ventricular dysfunction[45]. BAG3 serves a range of important functions in myocytes via its multiple protein-binding domains. BAG3 contributes to protein homeostasis through interactions with HSP70 (HSPA1A), Hsc70 (HSPA8), HSPB1, dynein and the ubiquitin ligase STUB1 to mediate selective macroautophagy[46,47]. This particularly important in post-mitotic cells to ensure clearing of misfolded and degraded proteins that arise with aging and in the context of cell stress or injury[48]. Through the WW protein domain, BAG3 participates in chaperone-assisted selective autophagy (CASA), of which the actin-cross linking protein filamin has been implicated as a client, a process that is dependent upon an interaction with synaptopodin-2 (SYNPO2), a paralogue of the *SYNPO2L* gene described above[49].

***ATXN2*** **(rs4766578)**

The sentinel variant *at this locus is located in the intron of ATXN2. Variants in and around ATXN2 and the adjacent gene SH2B3* have been associated with multiple traits, including include white blood cell counts, body-mass index, coronary artery disease, diabetes and blood pressure (**Supplementary Tables 2 and 4**). Both *ATXN2* and *SH2B3* are ubiquitously expressed. *ATXN2 codes* for Ataxin-2, is a cytoplasmic protein with important signalling functions that modulates ribosomal translation, mitochondrial function and mTOR signalling[50]. It is ubiquitously expressed in many tissues. Deficiency of Ataxin-2 is associated with effects on several metabolic pathways and is associated with lipid metabolism, obesity, and diabetes[51,52]. *SH2B3* codes for a protein involved in hematopoiesis and inflammation[53].

***FTO*** **(rs56094641)**

Variants at the *FTO* locus were first found to be associated with BMI and risk of obesity in 2007[54], The HF sentinel variant at this locus is in high linkage disequilibrium with a known variant for BMI at this locus (rs1558902, $r^2$ 0.98)[55]. Loss of Fto in mice leads to reduced body weight and fat mass[56]. Recent studies have also implicated the adjacent gene *IRX3* as a functional transcription factor in relation to BMI. Variants within *FTO* were shown to form long-range interactions affecting *IRX3* expression. A reduction in body weight of 25 to 30% has been shown in *Irx3*-deficient mice, providing a direct link between *IRX3* expression and regulation of body mass and composition[57].

### 2. Description of participating studies

#### Atherosclerosis Risk in Communities Study (ARIC)[58]

The ARIC study is a population-based prospective cohort study of cardiovascular disease sponsored by the National Heart, Lung, and Blood Institute (NHLBI). ARIC included 15,792 individuals, predominantly European American and African American, aged 45-64 years at baseline (1987-89), chosen by probability sampling from four US communities. Cohort members completed three additional triennial follow-up examinations, a fifth exam in 2011-2013, and a sixth exam in 2016-2017.

#### A systems BIOlogy Study to TAilored Treatment in Chronic Heart Failure (BIOSTAT-CHF) – Validation cohort[59]

BIOSTAT-CHF was a multicentre, multinational, prospective, observational study. Thel validation cohort consisted of 1,738 patients recruited from 6 hospitals in Scotland from 2010-2014.

#### Cardiovascular Health Study (CHS)[60,61]

The Cardiovascular Health Study (CHS) is a population-based cohort study of risk factors for coronary heart disease and stroke in adults ≥65 years conducted across four field centers. The original predominantly European ancestry cohort of 5,201 persons was recruited in 1989-1990 from random samples of the Medicare eligibility lists; subsequently, an additional predominantly African-American cohort of 687 persons was enrolled in 1992-1993 for a total sample of 5,888. For this analysis, only participants with European ancestry were included. European ancestry participants were excluded from the GWAS study sample due to the presence at study baseline of coronary heart disease, congestive heart failure, peripheral vascular disease, valvular heart disease, stroke or transient ischemic attack or lack of available DNA.

**The Copenhagen Cardiovascular Genetic study (COGEN)**[62]

The Copenhagen Cardiovascular Genetic study (COGEN) is a biobank that has collected superfluous whole blood from patients admitted to six cardiology departments in the Capital region of Copenhagen from 2010-2017.

**deCODE Heart Failure Study (deCODE)**[63]

The deCODE Icelandic heart failure (HF) sample set included patients diagnosed with HF at Landspitali – The National University Hospital (LUH) in Reykjavik. The controls included population controls from the Icelandic genealogical database and individuals recruited through different genetic studies at deCODE genetics. Individuals of non-Icelandic origin were excluded from the study.

**DiscovEHR**[64]

DiscovEHR is a collaboration between Regeneron Genetics Center and Geisinger Health System. The population is derived from patients who have previously consented to participate in the Geisinger MyCode Community Health Initiative.

**Estonian Genome Center at the University of Tartu (EGCUT)**[65]

The Estonia biobank is a population-based cohort of the Estonian Genome Center at the University of Tartu (EGCUT), which was established in 2001. Subjects were recruited at random and represent about 5% of the Estonian population

**Eplerenone Post-Acute Myocardial Infarction Heart Failure Efficacy and Survival Study (EPHESUS)**[66]

The EPHESUS was a randomized, placebo-controlled clinical trial of eplerenone that enrolled a total of 6,642 patients in 37 countries between 1999 and 2001. Participants were randomized within 3-14 days following a documented acute myocardial infarction with left ventricular dysfunction, demonstrated by left ventricular ejection fraction ≤ 40% and clinical evidence of heart failure. Participants were followed up an average of 16 months. Eplerenone was developed by Pharmacia, which was later acquired by Pfizer, and eplerenone is marketed as Inspra. The controls (A9011027) were recruited in a multi-site, cross-sectional, non-treatment prospective trial to collect data, including DNA, from elderly subjects in the US who were cognitively normal and free of psychiatric diseases or clinically significant cardiovascular disease.

**The European Prospective Investigation of Cancer, Norfolk study (EPIC-Norfolk)**[67]

The EPIC-Norfolk study is a prospective population-based cohort study which recruited 25,639 men and women aged 40-79 years at baseline between 1993 and 1997 from 35 participating general practices in Norfolk, UK. Individuals attended for a baseline health check including the provision of blood samples for concurrent and future analysis. They provided consent to future linkage to medical record information and a wide range of follow-up studies for different disease endpoints (including incident T2DM) have subsequently been undertaken, and further health check visits have been conducted since the baseline visit (see www.srl.cam.ac.uk/epic).

**Estonian Genome Center at the University of Tartu (EGCUT)**[65]

The Estonia biobank is a population-based cohort of the Estonian Genome Center at the University of Tartu (EGCUT), which was established in 2001. Subjects were recruited at random and represent about 5% of the Estonian population.

**Framingham Heart Study (FHS)**[68–70]

The FHS is a community-based cohort that enrolled three generations of participants. The objective of the FHS was to identify common factors or characteristics that contribute to CVD by following its development over a long period of time in a large group of participants who had not yet developed overt symptoms of CVD or suffered a heart attack or stroke. The researchers recruited 5,209 men and women between the ages of 30 and 62 from the town of Framingham, Massachusetts, and began the first round of extensive physical examinations and lifestyle interviews that they would later analyze for common patterns related to CVD development. Since 1948, the subjects have continued to return to the study every two years for a detailed medical history, physical examination, and laboratory tests, and in 1971, the study enrolled a second generation - 5,124 of the original participants' adult children and their spouses - to participate in similar examinations. In 1994, the need to establish a new study reflecting a more diverse community of Framingham was recognized, and the first Omni cohort of the Framingham Heart Study was enrolled. In April 2002 the Study entered a new phase, the enrollment of a third generation of participants, the grandchildren of the Original Cohort. In 2003, a second group of Omni participants was enrolled.

**FINRISK**[71]

FINRISK is a series of health examination surveys carried out by the National Institute for Health and Welfare (formerly National Public Health Institute) of Finland every five years since 1972. The surveys are based on random population samples from five (or six in 2002) specified geographical areas of Finland. The samples have been stratified by 10-year age group, sex and study area. The sample sizes have varied from approximately from 7,000 to 13,000 individuals and the participation rates from 90% to 60% in different study years. The age-range was 25-64 years until 1992 and 25-74 since 1997. The survey included a self-administered questionnaire, a standardized clinical examination carried out by specifically trained study nurses and drawing of a blood sample. DNA has been collected since the 1992 survey from approximately 34,000 participants. The follow-up of FINRISK participants takes place with annual record linkage of FINRISK data to the country-wide electronic health care registers, the National Causes-of-Death Register, Hospital Discharge Register including ambulatory visits to specialist health care facilities, Drug Reimbursement Registers and the Cancer register. The record linkage is carried out using the personal ID code, which is unique to every permanent resident of Finland.

**Genetics of Diabetes Audit and Research Tayside Scotland (GoDARTS)**[72]

GoDARTS is a cohort study of 18,306 participants, 10,149 with type 2 diabetes and 8157 healthy controls at baseline recruited from 1996 to 2009 in Tayside, Scotland. Genetic data are available for 8,564 T2D cases and 4,586 controls. Overall, 53.33% of the cohort are male. The majority of the cohort are Caucasian (99.70%) and the median age at recruitment was 64 years. Patients consented to electronic health record linkage to allow follow-up on mortality, hospitalisations and investigations including echocardiography.

**The Genetic Risk Assessment of Defibrillator Events (GRADE)**

The GRADE study was designed to identify genetic modifiers of arrhythmic risk. Inclusion criteria were: patients who were ≥18 years of age with a diagnosis of at least moderate systolic left ventricular dysfunction (EF ≤30%), and who had an ICD at the University of Pittsburgh Medical Center, Emory University Medical Center, Massachusetts General Hospital, Ohio State University Medical Center, Mid-Ohio Cardiology or the Pittsburgh Veterans Affairs Medical Center. Subjects were excluded if they had intractable Class IV heart failure, and conditions (other than HF) that were expected to limit survival to less than 6 months. The institutional review boards of participating medical centers approved the study and each patient gave written informed consent prior to participation. This study was conducted according to the guidelines laid down in the Declaration of Helsinki and the trial was registered at www.clinicaltrials.gov (NCT 02045043). Controls were drawn from the Broad AF study, a large case-control cohort study for atrial fibrillation[73]. Unrelated individuals of genetically inferred European ancestry free of AF and HF were selected as controls.

**The LUdwigshafen RIsk and Cardiovascular Health (LURIC) study**[74]

The LURIC study is a monocentric hospital based prospective study including 3316 individuals referred for coronary angiography recruited in the Ludwigshafen Cardiac Center, southwestern Germany from 1997 – 2000. Clinical indications for angiography were chest pain or a positive non-invasive stress test suggestive of myocardial ischemia. To limit clinical heterogeneity, individuals suffering from acute illnesses other than acute coronary syndrome, chronic non-cardiac diseases and a history of malignancy within the five past years were excluded. All participants completed a detailed questionnaire which gathered information on medical history, clinical, and lifestyle factors. Fasting blood samples were obtained by venipuncture in the early morning and stored for later analyses.

**Malmö Diet and Cancer Study (MDCS)**[75]

MDCS is a community-based prospective cohort of middle-aged individuals from Southern Sweden. In total, 30,447 subjects attended a baseline exam in 1991-1996 when they filled out a questionnaire, underwent anthropometric measurements and donated peripheral venous blood samples. The study was approved by the local ethics committee and all participants provided written informed consent. Prevalent or incident cases of heart failure were ascertained from nation-wide hospital registers with high validity as previously reported[76]. Genotyping was performed in a nested case-cohort design including 8,346 subjects with complete data of which 755 cases with incident heart failure.

**Prospective Investigation of the Vasculature in Uppsala Seniors (PIVUS)**[77]

The PIVUS study, initiated in 2002, is a community-based prospective cohort comprising 1016 randomly selected men and women aged 70 years in Uppsala county. Subjects were re-investigated at age 75 and age 80 years. The PIVUS-study is unique in its detailed characterization of vascular and cardiac function and morphology. The cohort is also well characterized with regards to genomics, epigenomics, proteomics and metabolomics. Ten-year follow-up data on cardiovascular events and mortality is available.

**Prevention of REnal and Vascular ENd-stage Disease (PREVEND)**[78]

The PREVEND Study is a prospective, observational cohort study, focussed to assess the impact of elevated urinary albumin loss in non-diabetic subjects on future cardiovascular and renal disease. PREVEND is an acronym for Prevention of REnal and Vascular ENd-stage Disease. This study started with a population survey on the prevalence of micro-albuminuria and generation of a study cohort of the general population. The goal is to monitor this cohort for the long-term development of cardiac-, renal- and peripheral vascular end-stage disease. For that purpose, the participants receive questionnaires on events and are seen every three/four years for a survey on cardiac-, renal- and peripheral vascular morbidity. The population is formed of Groningen inhabitants aged 28 to 75 years, who agreed to give a morning urine sample and to answer a short questionnaire. Of the 85,421 subjects invited to participate, 40,856 responded. The final sample is consisted of 8,592 consenting subjects with a morning urinary albumin concentration (UAC) of >10 mg/L and an a-select sample of those with an UAC <10 mg/L who completed the first screening; half of which were genotyped.

**PROspective Study of Pravastatin in the Elderly at Risk for vascular disease (PROSPER)**[79–81]

PROSPER was a prospective multicenter randomized placebo-controlled trial to assess whether treatment with pravastatin diminishes the risk of major vascular events in elderly. Between December 1997 and May 1999, we screened and enrolled subjects in Scotland (Glasgow), Ireland (Cork), and the Netherlands (Leiden). Men and women aged 70-82 years were recruited if they had pre-existing vascular disease or increased risk of such disease because of smoking, hypertension, or diabetes. A total number of 5,804 subjects were randomly assigned to pravastatin or placebo. A large number of prospective tests were performed including Biobank tests and cognitive function measurements.

**Rotterdam Study 1 (Rotterdam 1)**[82]

The Rotterdam Study is a prospective population-based cohort study that addresses determinants and occurrence of cardiovascular, neurological, ophthalmologic, psychiatric, and endocrine diseases in the elderly. At present the Rotterdam Study incorporates three cohorts that were established in 1989, 2000, 2006, and 2015 respectively. In 1989 all residents of Ommoord, a suburb of Rotterdam, aged 55 years and over were invited to participate. A total of 7,983 out of 10,275 men and women entered the study (response rate 78 percent, click here for these baseline response data). Baseline data were collected from 1990 until 1993. From 2009 onwards we are seeing these participants for the fifth time. HF case ascertainment has been detailed elsewhere[83].

**Study of Health in Pomerania (SHIP)**[84]

The Study of Health in Pomerania (SHIP) is a population-based project in West Pomerania, the north-east area of Germany. A sample from the population aged 20 to 79 years was drawn from population registries. First, the three cities of the region (with 17,076 to 65,977 inhabitants) and the 12 towns (with 1,516 to 3,044 inhabitants) were selected, and then 17 out of 97 smaller towns (with less than 1,500 inhabitants), were drawn at random. Second, from each of the selected communities, subjects were drawn at random, proportional to the population size of each community and stratified by age and gender. Only individuals with German citizenship and main residency in the study area were included. Finally, 7,008 subjects were sampled, with 292 persons of each gender in each of the twelve five-year age strata. In order to minimize drop-outs by migration or death, subjects were selected in two waves. The net sample (without migrated or deceased persons) comprised 6,267 eligible subjects. Selected persons received a maximum of three written invitations. In case of non-response, letters were followed by a phone call or by home visits if contact by phone was not possible. The SHIP population finally comprised 4,308 participants (corresponding to a final response of 68.8%). Non-fasting blood samples were drawn from the cubital vein in the supine position. The samples were taken between 07:00 AM and 04:00 PM, and serum aliquots were prepared for immediate analysis and for storage at -80 °C in the Integrated Research Biobank (Liconic, Liechtenstein).

**Stabilization of Plaque using Darapladib-Thrombolysis in Myocardial Infarction 52 (SOLID)**[85]

The SOLID-TIMI 52 was a multinational, double-blind trial that enrolled 13,026 participants who had been hospitalized with an acute coronary syndrome event in the past 30 days and randomized them to once daily darapladib or placebo for a median follow-up of 2.5 years (7). The primary endpoint was CHD death, MI or urgent coronary revascularization. In each country, the study was approved by national regulatory authorities and by local ethics committees or institutional review boards, according to local regulations. This study did not meet its primary endpoint.

**TwinGene (TwinGene)**[86]

TwinGene is a population-based cohort within the Swedish Twin Registry. TwinGene participants were born between 1911 and 1958, they had previously participated in the Screening Across the Lifespan Twin (SALT) study, a computer-assisted telephone interview conducted between 1998 and 2002. During 2004 and 2008, ~12000 TwinGene participants donated their blood at the local health care facility after overnight fasting, and their height, weight, hip and waist circumference, as well as their blood pressures were measured. The zygosity was identified by self-reported childhood resemblance and DNA markers. Both twins within a pair had to be alive and consent for future participation.

**UK Biobank**[87,88]

The UK Biobank is a large, population-based prospective cohort with extensive genetic and phenotypic data collected on approximately 500,000 individuals aged 40–69 years recruited from across the UK between 2006 and 2010. Collected information include socio-demographics, lifestyle, and health-related factors, physical measures, biological samples (blood, urine, and saliva) for genomics and biochemical markers assessments, linked electronic health records, disease registers, and death register, with a planned repeat assessments and multi-modal imaging. The UK Biobank genetic data contains genotypes for 488,377 participants assayed using two very similar genotyping arrays with extensive phasing and genotype imputation[87,88].

**Uppsala Longitudinal Study of Adult Men (ULSAM)**[89,90]

The ULSAM study is a longitudinal community based cohort study of 50 year old men that started in 1970 (n=2322). All 50 year old men in Uppsala were invited to participate. Subjects were reinvestigated at the ages of 60, 70, 77, 82 and 88 years. GWAS data is available in approximately half of the study sample. Untargeted metabolomics and proteomics data is available both in serum and urine in subsamples of the cohort. A large number of cardiovascular events and mortality data is available with more than 20 years of follow-up.

**Penn Heart Failure Study (PHFS)**[91,92]

A case-control genome-wide association study comparing patients with prevalent heart failure referred for evaluation and treatment at a heart failure specialty centre. Prevalent heart failure was identified by a heart failure cardiologist based on clinical evaluation and cardiac imaging.

**Women's Genome Health Study (WGHS)**[93]

The Women's Genome Health Study (WGHS) is a prospective cohort of initially healthy, female North American health care professionals at least 45 years old at baseline representing participants in the Women's Health Study (WHS) who provided a blood sample at baseline and consent for blood-based analyses. The WHS was a 2x2 trial beginning in 1992-1994 of vitamin E and low dose aspirin in prevention of cancer and cardiovascular disease with about 10 years of follow-up. Since the end of the trial, follow-up has continued in observational mode. Additional information related to health and lifestyle were collected by questionnaire throughout the WHS trial and continuing observational follow-up. WGHS genetic data are currently not publicly accessible. Genetic analysis in the WGHS has been approved by the IRB of Brigham and Women's Hospital.

### 3. Description of participating consortia

#### Myocardial Applied Genomics Network (MAGNet)

The MAGNet repository (http://www.med.upenn.edu/magnet/) includes samples from normal donors and from patients with heart failure at the time of cardiac transplantation. The study protocol was approved by the Institutional Review Board at the University of Pennsylvania, and all patients provided written informed consent to participate.

### 4. Acknowledgements

**Atherosclerosis Risk in Communities Study (ARIC)**

The Atherosclerosis Risk in Communities Study is carried out as a collaborative study supported by National Heart, Lung, and Blood Institute contracts N01-HC-55015, N01-HC-55016, N01-HC-55018, N01-HC-55019, N01-HC-55020, N01-HC55021, N01-HC-55022, R01HL087641, R01HL59367, R01HL086694 and RC2 HL102419; National Human Genome Research Institute contract U01HG004402; and National Institutes of Health contract HHSN268200625226C. The authors thank the staff and participants of the ARIC study for their important contributions. Infrastructure was partly supported by Grant Number UL1RR025005, a component of the National Institutes of Health and NIH Roadmap for Medical Research.

**A systems BIOlogy Study to TAilored Treatment in Chronic Heart Failure (BIOSTAT-CHF)**

This project was funded by a grant from the European Commission (FP7-242209-BIOSTAT-CHF; EudraCT 2010–020808–29).

**Cardiovascular Health Study (CHS)**

This CHS research was supported by NHLBI contracts HHSN268201200036C, HHSN268200800007C, HHSN268201800001C, HHSN268200960009C, N01HC55222, N01HC85079, N01HC85080, N01HC85081, N01HC85082, N01HC85083, N01HC85086; and NHLBI grants U01HL080295, R01HL087652, R01HL105756, R01HL103612, R01HL120393, and U01HL130114 with additional contribution from the National Institute of Neurological Disorders and Stroke (NINDS). Additional support was provided through R01AG023629 from the National Institute on Aging (NIA). A full list of principal CHS investigators and institutions can be found at CHS-NHLBI.org. The provision of genotyping data was supported in part by the National Center for Advancing Translational Sciences, CTSI grant UL1TR001881, and the National Institute of Diabetes and Digestive and Kidney Disease Diabetes Research Center (DRC) grant DK063491 to the Southern California Diabetes Endocrinology Research Center.

**deCODE Heart Failure Study (deCODE)**

We at deCODE thank the women and men of Iceland that have participated in our studies and our colleagues that contributed to data collection and processing.

**DiscovEHR**

We acknowledge and thank all participants in Geisinger's MyCode Community Health Initiative for their support and permission to use their health and genomic information in the DiscovEHR collaboration. This work was supported by the Regeneron Genetics Center and Geisinger.

**Estonian Genome Center at the University of Tartu (EGCUT)**

This study was supported by Estonian Research Council Grant IUT20-60, EU, H2020 grant 692145, European Union through the European Regional Development Fund (Project No. 2014-2020.4.01.15-0012) GENTRANSMED.

**Eplerenone Post-Acute Myocardial Infarction Heart Failure Efficacy and Survival Study (EPHESUS)**

The EPHESUS was supported by Pfizer, Inc.

**The European Prospective Investigation of Cancer, Norfolk study (EPIC-Norfolk)**

The EPIC-Norfolk Study is supported by programme grants from the Medical Research Council UK (G1000143) and Cancer Research UK (C864/A14136) and with additional support from the European Union, Stroke Association, British Heart Foundation, Research into Ageing, Department of Health, The Wellcome Trust and the Food Standards Agency. NJW and CL also acknowledge support from the Medical Research Council, UK (MC_UU_12015/1; MC_PC_13048). We thank all EPIC participants and staff for their contribution to the study, and thank staff from the Technical, Field Epidemiology and Data Functional Group Teams of the Medical Research Council Epidemiology Unit in Cambridge, UK, for carrying out sample preparation, DNA provision and quality control, genotyping and data handling work.

**Framingham Heart Study (FHS)**

This work was conducted using data and resources from the Framingham Heart Study (FHS) of the National Heart Lung and Blood Institute and Boston University School of Medicine. The study was supported by the National Heart, Lung and Blood Institute's Framingham Heart Study (Contract No. N01-HC-25195 and HHSN268201500001I) and its contract with Affymetrix, Inc for genotyping services (Contract No.N02-HL-6-4278). The work was also supported by R01 HL093328, R01 HL105993, and R01 HL71039 (PI: Ramachandran).

**FINRISK**

V.S. has been supported by the Finnish Foundation for Cardiovascular Research.

**Genetics of Diabetes Audit and Research Tayside Scotland GoDARTS)**

The Wellcome Trust United Kingdom Type 2 Diabetes Case Control Collection (supporting GoDARTS) was funded by the Wellcome Trust (072960/Z/03/Z, 084726/Z/08/Z, 084727/Z/08/Z, 085475/Z/08/Z, 085475/B/08/Z) and as part of the EU IMI-SUMMIT programme. We acknowledge the support of the Health Informatics Centre, University of Dundee, for managing and supplying the anonymized data and NHS Tayside, the original data owner.

**The Genetic Risk Assessment of Defibrillator Events (GRADE)**

NIH-NHLBI R01 HL77398 (Genetic Modulators of Sudden Death). S.L. is supported by NIH grant 1R01HL139731 and American Heart Association 18SFRN34250007.

**The LUdwigshafen RIsk and Cardiovascular Health (LURIC) study**

We extend our appreciation to the participants of the LURIC study; without their collaboration, this article would not have been written. We thank the LURIC study team who were either temporarily or permanently involved in patient recruitment as well as sample and data handling, in addition to the laboratory staff at the Ludwigshafen General Hospital and the Universities of Freiburg and Ulm, Germany. LURIC has received funding from the 7th Framework Program (RiskyCAD, grant agreement number 305739 and Atheroremo, grant agreement number 201668) of the European Union.

**Malmö Diet and Cancer Study (MDCS)**

J. Gustav Smith was supported by grants from the Swedish Heart-Lung Foundation (2016-0134 and 2016-0315), the Swedish Research Council (2017-02554), the European Research Council (ERC-STG-2015-679242), the Crafoord Foundation, Skåne University Hospital, the Scania county, governmental funding of clinical research within the Swedish National Health Service, a generous donation from the Knut and Alice Wallenberg foundation to the Wallenberg Center for Molecular Medicine in Lund, and funding from the Swedish Research Council (Linnaeus grant Dnr 349-2006-237, Strategic Research Area Exodiab Dnr 2009-1039) and Swedish Foundation for Strategic Research (Dnr IRC15-0067) to the Lund University Diabetes Center. The Malmo Diet and Cancer Study was made possible by grants from the Swedish Cancer Society, the Swedish Medical Research Council, the Swedish Dairy Association, and the Malmo city council.

**Penn Heart Failure Study (PHFS)**

The study was supported by NIH grants (NIH R01L088577 and NIH R01H105993).

**Prevention of REnal and Vascular ENd-stage Disease (PREVEND)**

The Prevention of Renal and Vascular Endstage Disease Study (PREVEND) genetics is supported by the Dutch Kidney Foundation (Grant E033), the EU project grant GENECURE (FP-6 LSHM CT 2006 037697), the National Institutes of Health (grant LM010098), the Netherlands organisation for health research and development (NWO VENI grant 916.761.70), and the Dutch Inter University Cardiology Institute Netherlands (ICIN). Niek Verweij was supported by NWO VENI grant 016.186.125.

**PROspective Study of Pravastatin in the Elderly at Risk for vascular disease (PROSPER)**

The PROSPER study was supported by an investigator-initiated grant obtained from Bristol-Myers Squibb. Support for genotyping was provided by the seventh framework program of the European commission (grant 223004) and by the Netherlands Genomics Initiative (Netherlands Consortium for Healthy Aging grant 050-060-810). J.W.J. is an Established Clinical Investigator of the Netherlands Heart Foundation (grant 2001 D 032).

**Study of Health in Pomerania (SHIP)**

SHIP is part of the Community Medicine Research net of the University of Greifswald, Germany, which is funded by the Federal Ministry of Education and Research (grants no. 01ZZ9603, 01ZZ0103, and 01ZZ0403), the Ministry of Cultural Affairs as well as the Social Ministry of the Federal State of Mecklenburg-West Pomerania, and the network 'Greifswald Approach to Individualized Medicine (GANI_MED)' funded by the Federal Ministry of Education and Research (grant 03IS2061A). Genome-wide data have been supported by the Federal Ministry of Education and Research (grant no. 03ZIK012) and a joint grant from Siemens Healthineers, Erlangen, Germany and the Federal State of Mecklenburg- West Pomerania. The University of Greifswald is a member of the Caché Campus program of the InterSystems GmbH.

**Stabilization of Plaque using Darapladib-Thrombolysis in Myocardial Infarction 52 (SOLID)**

SOLID-TIMI 52 was funded by GlaxoSmithKline.

**TwinGene (TwinGene)**

TwinGene received funding from the Swedish Research Council (M-2005-1112), GenomEUtwin (EU/QLRT-2001-01254; QLG2-CT-2002-01254), NIH DK U01-066134, The Swedish Foundation for Strategic Research (SSF) and the Heart and Lung foundation no. 20070481. TwinGene is part of the Swedish Twin Registry which is managed by Karolinska Institutet and receives funding through the Swedish Research Council (2017–00641).

**UK Biobank (UKBiobank)**

This work was supported in part by grants to R.T.L. from the EU/EFPIA Innovative Medicines Initiative 2 Joint Undertaking BigData@Heart grant no. 116074, MRC Proximity to Discovery Award Scheme, the American Heart Association Institute for Precision Mecidine, Pfizer Ltd, and was supported by the National Institute for Health Research University College London Hospitals Biomedical Research Centre. A. H. is supported by the British Heart Foundation Cardiovascular Biomedicine PhD studentship. R.T.L is supported by a UK Research and Innovation Rutherford Fellowship and was previously supported by a National Institutes of Health Research Clinical Lectureship.

**Uppsala Longitudinal Study of Adult Men (ULSAM)**

J.Ä. is supported by the Swedish Research Council and the Swedish Heart Lung foundation. C.M.L is supported by the Li Ka Shing Foundation, WT-SSI/John Fell funds and by the NIHR Biomedical Research Centre, Oxford, by Widenlife and NIH (5P50HD028138-27).

**Women's Genome Health Study (WGHS)**

The WGHS is supported by the National Heart, Lung, and Blood Institute (HL043851 and HL080467, HL099355) and the National Cancer Institute (CA047988 and UM1CA182913), with collaborative scientific support and funding for genotyping provided by Amgen.

### Supplementary References


1. Willer, C. J. *et al.* Discovery and refinement of loci associated with lipid levels. *Nat. Genet.* **45**, 1274–1283 (2013).

2. Samani, N. J. *et al.* Genomewide Association Analysis of Coronary Artery Disease. *N. Engl. J. Med.* **357**, 443–453 (2007).

3. Nikpay, M. *et al.* A comprehensive 1,000 Genomes-based genome-wide association meta-analysis of coronary artery disease. *Nat. Genet.* **47**, 1121–1130 (2015).

4. Jang, C.-Y. *et al.* DDA3 recruits microtubule depolymerase Kif2a to spindle poles and controls spindle dynamics and mitotic chromosome movement. *J. Cell Biol.* **181**, 255–67 (2008).

5. Nurnberg, S. T. *et al.* From Loci to Biology: Functional Genomics of Genome-Wide Association for Coronary Disease. *Circ. Res.* **118**, 586–606 (2016).

6. Tönjes, A. *et al.* Genome-wide meta-analysis identifies novel determinants of circulating serum progranulin. *Hum. Mol. Genet.* **27**, 546–558 (2018).

7. Ye, J. *et al.* A Functional Variant Associated with Atrial Fibrillation Regulates PITX2c Expression through TFAP2a. *Am. J. Hum. Genet.* **99**, 1281–1291 (2016).

8. Tao, G. *et al.* Pitx2 promotes heart repair by activating the antioxidant response after cardiac injury. *Nature* **534**, 119–123 (2016).

9. Mori, Y. *et al.* Involvement of selective autophagy mediated by p62/SQSTM1 in KLHL3-dependent WNK4 degradation. *Biochem. J.* **472**, 33–41 (2015).

10. Shibata, S., Zhang, J., Puthumana, J., Stone, K. L. & Lifton, R. P. Kelch-like 3 and Cullin 3 regulate electrolyte homeostasis via ubiquitination and degradation of WNK4. *Proc. Natl. Acad. Sci. U. S. A.* **110**, 7838–43 (2013).

11. Louis-Dit-Picard, H. *et al.* KLHL3 mutations cause familial hyperkalemic hypertension by impairing ion transport in the distal nephron. *Nat. Genet.* **44**, 456–460 (2012).

12. Boyden, L. M. *et al.* Mutations in kelch-like 3 and cullin 3 cause hypertension and electrolyte abnormalities. *Nature* **482**, 98–102 (2012).

13. Harper, J. W., Adami, G. R., Wei, N., Keyomarsi, K. & Elledge, S. J. The p21 Cdk-interacting protein Cip1 is a potent inhibitor of G1 cyclin-dependent kinases. *Cell* **75**, 805–16 (1993).

14. Campisi, J. Aging, cellular senescence, and cancer. *Annu. Rev. Physiol.* **75**, 685–705 (2013).

15. Tane, S. *et al.* CDK inhibitors, p21Cip1 and p27Kip1, participate in cell cycle exit of mammalian cardiomyocytes. *Biochem. Biophys. Res. Commun.* **443**, 1105–1109 (2014).

16. Zhang, D. *et al.* REST regulates the cell cycle for cardiac development and regeneration. *Nat. Commun.* **8**, 1979 (2017).

17. Bedelbaeva, K. *et al.* Lack of p21 expression links cell cycle control and appendage regeneration in mice. *Proc. Natl. Acad. Sci. U. S. A.* **107**, 5845–50 (2010).

18. Mattioli, E. *et al.* Altered modulation of lamin A/C-HDAC2 interaction and *p21* expression during oxidative stress response in HGPS. *Aging Cell* **17**, e12824 (2018).

19. Mattioli, E. *et al.* Altered modulation of lamin A/C-HDAC2 interaction and *p21* expression during oxidative stress response in HGPS. *Aging Cell* e12824 (2018). doi:10.1111/acel.12824

20. Clarke, R. *et al.* Genetic Variants Associated with Lp(a) Lipoprotein Level and Coronary Disease. *N. Engl. J. Med.* **361**, 2518–2528 (2009).

21. Trégouët, D.-A. *et al.* Genome-wide haplotype association study identifies the SLC22A3-LPAL2-LPA gene cluster as a risk locus for coronary artery disease. *Nat. Genet.* **41**, 283–285 (2009).

22. Austin, M. A. *et al.* Lipoprotein(a) in women twins: heritability and relationship to

apolipoprotein(a) phenotypes. *Am. J. Hum. Genet.* **51**, 829–40 (1992).

23. Burgess, S. *et al.* Association of LPA Variants With Risk of Coronary Disease and the Implications for Lipoprotein(a)-Lowering Therapies: A Mendelian Randomization Analysis. *JAMA Cardiol.* **3**, 619–627 (2018).

24. Arimany-Nardi, C., Koepsell, H. & Pastor-Anglada, M. Role of SLC22A1 polymorphic variants in drug disposition, therapeutic responses, and drug-drug interactions. *Pharmacogenomics J.* **15**, 473–87 (2015).

25. Fan, M. *et al.* Two Chromosome 9p21 Haplotype Blocks Distinguish Between Coronary Artery Disease and Myocardial Infarction Risk. *Circ. Cardiovasc. Genet.* **6**, 372–380 (2013).

26. Holdt, L. M. *et al.* ANRIL expression is associated with atherosclerosis risk at chromosome 9p21. *Arterioscler. Thromb. Vasc. Biol.* **30**, 620–7 (2010).

27. Zhang, H., Mooney, C. J. & Reilly, M. P. ABO Blood Groups and Cardiovascular Diseases. *Int. J. Vasc. Med.* **2012**, 641917 (2012).

28. von Beckerath, N. *et al.* ABO locus O1 allele and risk of myocardial infarction. *Blood Coagul. Fibrinolysis* **15**, 61–7 (2004).

29. van Eldik, W. *et al.* Z-disc protein CHAPb induces cardiomyopathy and contractile dysfunction in the postnatal heart. *PLoS One* **12**, e0189139 (2017).

30. Beqqali, A. *et al.* CHAP is a newly identified Z-disc protein essential for heart and skeletal muscle function. *J. Cell Sci.* **123**, 1141–50 (2010).

31. Frey, N. *et al.* Calsarcin-2 deficiency increases exercise capacity in mice through calcineurin/NFAT activation. *J. Clin. Invest.* **118**, 3598–608 (2008).

32. Faulkner, G. *et al.* FATZ, a filamin-, actinin-, and telethonin-binding protein of the Z-disc of skeletal muscle. *J. Biol. Chem.* **275**, 41234–42 (2000).

33. Takada, F. *et al.* Myozenin: an alpha-actinin- and gamma-filamin-binding protein of skeletal muscle Z lines. *Proc. Natl. Acad. Sci. U. S. A.* **98**, 1595–600 (2001).

34. Arola, A. M. *et al.* Mutations in PDLIM3 and MYOZ1 encoding myocyte Z line proteins are infrequently found in idiopathic dilated cardiomyopathy. *Mol. Genet. Metab.* **90**, 435–40 (2007).

35. Osio, A. *et al.* Myozenin 2 is a novel gene for human hypertrophic cardiomyopathy. *Circ. Res.* **100**, 766–8 (2007).

36. Homma, S. *et al.* BAG3 deficiency results in fulminant myopathy and early lethality. *Am. J. Pathol.* **169**, 761–73 (2006).

37. Norton, N. *et al.* Genome-wide studies of copy number variation and exome sequencing identify rare variants in BAG3 as a cause of dilated cardiomyopathy. *Am. J. Hum. Genet.* **88**, 273–82 (2011).

38. Toro, R. *et al.* Familial Dilated Cardiomyopathy Caused by a Novel Frameshift in the BAG3 Gene. *PLoS One* **11**, e0158730 (2016).

39. Chami, N. *et al.* Nonsense mutations in BAG3 are associated with early-onset dilated cardiomyopathy in French Canadians. *Can. J. Cardiol.* **30**, 1655–61 (2014).

40. Esslinger, U. *et al.* Exome-wide association study reveals novel susceptibility genes to sporadic dilated cardiomyopathy. *PLoS One* **12**, e0172995 (2017).

41. Selcen, D. *et al.* Mutation in BAG3 causes severe dominant childhood muscular dystrophy. *Ann. Neurol.* **65**, 83–89 (2008).

42. Villard, E. *et al.* A genome-wide association study identifies two loci associated with heart failure due to dilated cardiomyopathy. *Eur. Heart J.* **32**, 1065–76 (2011).

43. Feldman, A. M. *et al.* Decreased levels of BAG3 in a family with a rare variant and in

idiopathic dilated cardiomyopathy. *J. Cell. Physiol.* **229**, 1697–702 (2014).

44. Franceschelli, S. *et al.* Bag3 gene expression is regulated by heat shock factor 1. *J. Cell. Physiol.* **215**, 575–577 (2008).

45. Myers, V. D. *et al.* Haplo-insufficiency of Bcl2-associated athanogene 3 in mice results in progressive left ventricular dysfunction, β-adrenergic insensitivity, and increased apoptosis. *J. Cell. Physiol.* **233**, 6319–6326 (2018).

46. Behl, C. BAG3 and friends: co-chaperones in selective autophagy during aging and disease. *Autophagy* **7**, 795–8 (2011).

47. Carra, S., Seguin, S. J., Lambert, H. & Landry, J. HspB8 chaperone activity toward poly(Q)-containing proteins depends on its association with Bag3, a stimulator of macroautophagy. *J. Biol. Chem.* **283**, 1437–44 (2008).

48. Behl, C. Breaking BAG: The Co-Chaperone BAG3 in Health and Disease. *Trends Pharmacol. Sci.* **37**, 672–688 (2016).

49. Ulbricht, A. *et al.* Cellular Mechanotransduction Relies on Tension-Induced and Chaperone-Assisted Autophagy. *Curr. Biol.* **23**, 430–435 (2013).

50. Aragam, K. G. *et al.* Phenotypic Refinement of Heart Failure in a National Biobank Facilitates Genetic Discovery. *Circulation* **139**, 489–501 (2019).

51. Kiehl, T.-R. *et al.* Generation and characterization of Sca2 (ataxin-2) knockout mice. *Biochem. Biophys. Res. Commun.* **339**, 17–24 (2006).

52. Lastres-Becker, I. *et al.* Insulin receptor and lipid metabolism pathology in ataxin-2 knock-out mice. *Hum. Mol. Genet.* **17**, 1465–1481 (2008).

53. Devallière, J. & Charreau, B. The adaptor Lnk (SH2B3): An emerging regulator in vascular cells and a link between immune and inflammatory signaling. *Biochem. Pharmacol.* **82**,

1391–1402 (2011).

54. Frayling, T. M. *et al.* A Common Variant in the FTO Gene Is Associated with Body Mass Index and Predisposes to Childhood and Adult Obesity. *Science (80-. ).* **316**, 889–894 (2007).

55. Locke, A. E. *et al.* Genetic studies of body mass index yield new insights for obesity biology. *Nature* **518**, 197–206 (2015).

56. Fischer, J. *et al.* Inactivation of the Fto gene protects from obesity. *Nature* **458**, 894–8 (2009).

57. Smemo, S. *et al.* Obesity-associated variants within FTO form long-range functional connections with IRX3. *Nature* **507**, 371–5 (2014).

58. The ARIC investigators. The Atherosclerosis Risk in Communities (ARIC) Study: design and objectives. *Am. J. Epidemiol.* (1989). doi:10.1093/oxfordjournals.aje.a115184

59. Voors, A. A. *et al.* A systems BIOlogy Study to TAilored Treatment in Chronic Heart Failure: rationale, design, and baseline characteristics of BIOSTAT-CHF. *Eur. J. Heart Fail.* (2016). doi:10.1002/ejhf.531

60. Fried, L. P. *et al.* The Cardiovascular Health Study: design and rationale. *Ann. Epidemiol.* **1**, 263–76 (1991).

61. Ives, D. G. *et al.* Surveillance and ascertainment of cardiovascular events. The Cardiovascular Health Study. *Ann. Epidemiol.* **5**, 278–85 (1995).

62. Hägg, S. *et al.* Adiposity as a cause of cardiovascular disease: a Mendelian randomization study. *Int. J. Epidemiol.* **44**, 578–86 (2015).

63. Gudbjartsson, D. F. *et al.* Large-scale whole-genome sequencing of the Icelandic population. *Nat. Genet.* **47**, 435–444 (2015).

64. Dewey, F. E. *et al.* Distribution and clinical impact of functional variants in 50,726 whole-

exome sequences from the DiscovEHR study. *Science* **354**, aaf6814 (2016).

65. Leitsalu, L. *et al.* Cohort Profile: Estonian Biobank of the Estonian Genome Center, University of Tartu. *Int. J. Epidemiol.* **44**, 1137–1147 (2015).

66. Pitt, B. *et al.* Eplerenone, a Selective Aldosterone Blocker, in Patients with Left Ventricular Dysfunction after Myocardial Infarction. *N. Engl. J. Med.* **348**, 1309–1321 (2003).

67. Day, N. *et al.* EPIC-Norfolk: study design and characteristics of the cohort. European Prospective Investigation of Cancer. *Br. J. Cancer* **80 Suppl 1**, 95–103 (1999).

68. DAWBER, T. R., MEADORS, G. F. & MOORE, F. E. Epidemiological approaches to heart disease: the Framingham Study. *Am. J. Public Health Nations. Health* **41**, 279–81 (1951).

69. Feinleib, M., Kannel, W. B., Garrison, R. J., McNamara, P. M. & Castelli, W. P. The Framingham Offspring Study. Design and preliminary data. *Prev. Med. (Baltim).* **4**, 518–25 (1975).

70. Splansky, G. L. *et al.* The Third Generation Cohort of the National Heart, Lung, and Blood Institute's Framingham Heart Study: design, recruitment, and initial examination. *Am. J. Epidemiol.* **165**, 1328–35 (2007).

71. Borodulin, K. *et al.* Cohort Profile: The National FINRISK Study. *Int. J. Epidemiol.* **47**, 696-696i (2018).

72. Hébert, H. L. *et al.* Cohort Profile: Genetics of Diabetes Audit and Research in Tayside Scotland (GoDARTS). *Int. J. Epidemiol.* **47**, 380-381j (2018).

73. Roselli, C. *et al.* Multi-ethnic genome-wide association study for atrial fibrillation. *Nat. Genet.* **50**, 1225–1233 (2018).

74. Winkelmann, B. R. *et al.* Rationale and design of the LURIC study--a resource for functional genomics, pharmacogenomics and long-term prognosis of cardiovascular disease.

*Pharmacogenomics* **2**, S1-73 (2001).

75. Smith, J. G., Platonov, P. G., Hedblad, B., Engström, G. & Melander, O. Atrial Fibrillation in the Malmö Diet and Cancer Study: A Study of Occurrence, Risk Factors and Diagnostic Validity. *European Journal of Epidemiology* **25**, 95–102 (2010).

76. Smith, J. G. *et al.* Assessment of Conventional Cardiovascular Risk Factors and Multiple Biomarkers for the Prediction of Incident Heart Failure and Atrial Fibrillation. *J. Am. Coll. Cardiol.* **56**, 1712–1719 (2010).

77. Lind, L., Fors, N., Hall, J., Marttala, K. & Stenborg, A. A comparison of three different methods to evaluate endothelium-dependent vasodilation in the elderly: the Prospective Investigation of the Vasculature in Uppsala Seniors (PIVUS) study. *Arterioscler. Thromb. Vasc. Biol.* **25**, 2368–75 (2005).

78. Pinto-Sietsma, S. J. *et al.* Urinary albumin excretion is associated with renal functional abnormalities in a nondiabetic population. *J. Am. Soc. Nephrol.* **11**, 1882–8 (2000).

79. Shepherd, J. *et al.* The design of a prospective study of Pravastatin in the Elderly at Risk (PROSPER). PROSPER Study Group. PROspective Study of Pravastatin in the Elderly at Risk. *Am. J. Cardiol.* **84**, 1192–7 (1999).

80. Shepherd, J. *et al.* Pravastatin in elderly individuals at risk of vascular disease (PROSPER): a randomised controlled trial. *Lancet (London, England)* **360**, 1623–30 (2002).

81. Trompet, S. *et al.* Replication of LDL GWAs hits in PROSPER/PHASE as validation for future (pharmaco)genetic analyses. *BMC Med. Genet.* **12**, 131 (2011).

82. Ikram, M. A. *et al.* The Rotterdam Study: 2018 update on objectives, design and main results. *Eur. J. Epidemiol.* **32**, 807–850 (2017).

83. Alberts, V. P. *et al.* Heart failure and the risk of stroke: the Rotterdam Study. *Eur. J. Epidemiol.* **25**, 807–12 (2010).

84. Volzke, H. *et al.* Cohort Profile: The Study of Health in Pomerania. *Int. J. Epidemiol.* **40**, 294–307 (2011).

85. O'Donoghue, M. L. *et al.* Effect of Darapladib on Major Coronary Events After an Acute Coronary Syndrome. *JAMA* **312**, 1006 (2014).

86. Magnusson, P. K. E. *et al.* The Swedish Twin Registry: Establishment of a Biobank and Other Recent Developments. *Twin Res. Hum. Genet.* **16**, 317–329 (2013).

87. Sudlow, C. *et al.* UK Biobank: An Open Access Resource for Identifying the Causes of a Wide Range of Complex Diseases of Middle and Old Age. *PLOS Med.* **12**, e1001779 (2015).

88. Bycroft, C. *et al.* The UK Biobank resource with deep phenotyping and genomic data. *Nature* **562**, 203–209 (2018).

89. Hedstrand, H. A study of middle-aged men with particular reference to risk factors for cardiovascular disease. *Ups. J. Med. Sci. Suppl.* **19**, 1–61 (1975).

90. Song, C. *et al.* Genetic Variants from Lipid-Related Pathways and Risk for Incident Myocardial Infarction. *PLoS One* **8**, e60454 (2013).

91. Cappola, T. P. *et al.* Loss-of-function DNA sequence variant in the CLCNKA chloride channel implicates the cardio-renal axis in interindividual heart failure risk variation. *Proc. Natl. Acad. Sci. U. S. A.* **108**, 2456–61 (2011).

92. Hu, R. *et al.* Genetic Reduction in Left Ventricular Protein Kinase C-α and Adverse Ventricular Remodeling in Human Subjects. *Circ. Genomic Precis. Med.* **11**, e001901 (2018).

93. Ridker, P. M. *et al.* Rationale, design, and methodology of the Women's Genome Health Study: a genome-wide association study of more than 25,000 initially healthy american women. *Clin. Chem.* **54**, 249–55 (2008).
